# Exogenous myristate fuels the growth of symbiotic arbuscular mycorrhizal fungi but disrupts their carbon-phosphorus exchange with host plants

**DOI:** 10.1101/2024.04.26.591230

**Authors:** Hanwen Chen, Tian Xiong, Baoxing Guan, Jiaqi Huang, Danrui Zhao, Yao Chen, Haoran Liang, Yingwei Li, Jingwen Wu, Shaoping Ye, Ting Li, Wensheng Shu, Jin-tian Li, Yutao Wang

## Abstract

Arbuscular mycorrhizal fungi (AMF) are obligate biotrophs that rely on symbiotic carbohydrates and, in particular, lipids derived from their host plants. However, it remains unclear whether symbiotic AMF can access exogenous non-symbiotic lipids in the presence of plant-derived carbon, complicating our understanding of their relationship with host plants. Here, we investigated the direct uptake of exogenous ^13^C_1_-labeled myristate by three symbiotic AMF species (*Rhizophagus irregularis*, *R. intraradices*, and *R. diaphanous*) and assessed their growth responses using AMF-carrot hairy root co-culture systems. Furthermore, we explored the environmental distribution of myristate, and evaluated the impact of exogenous myristate on the carbon-phosphorus exchange between *R. irregularis* and alfalfa or rice in a greenhouse experiment. Our results showed that symbiotic AMF can absorb exogenous myristate, as evidenced by ^13^C enrichment and transcriptional activation of fatty acid transport and metabolism genes in AMF extraradical hyphae. Myristate is commonly present in various soil and plant environments, and its application increased both intraradical and extraradical fungal biomass, possibly linked to suppressed mycorrhizal-activated defense responses in host roots. Unexpectedly, exogenous myristate reduced the mycorrhizal phosphorus benefits for both alfalfa and rice and decreased their symbiotic carbon allocation to root-colonizing AMF, suggesting that the application of exogenous myristate may not be a promising strategy to enhance AM symbiosis. These findings provide new insights into understanding and manipulating the nutritional interactions between AMF and host plants.

## INTRODUCTION

Arbuscular mycorrhizal fungi (AMF) are among the most widely distributed soil microorganisms, forming symbiotic relationships with approximately 70% of terrestrial vascular plants (Brundrett and Tedersoo, 2018). In this symbiosis, referred to as arbuscular mycorrhiza (AM), AMF develop an extensive extraradical hyphal network in the soil, which facilitates nutrient uptake for host plants—particularly phosphorus (P)—and enhances plant resilience to both biotic and abiotic stressors (Smith and Read, 2008). In return, host plants allocate 4−20% of their photosynthetically fixed carbon (C) to their fungal partners in the form of sugars and fatty acids (Helber et al., 2011; Jiang et al., 2017). This symbiotic C-P exchange, the cornerstone of AM symbiosis (Smith and Read, 2008), not only plays a pivotal role in the biogeochemical cycling of C and P elements but also acts as a key mechanism through which AMF influence plant community structure and productivity (Johnson, 2010; Wipf et al., 2019).

As one of the most ancient symbiotic relationships, AM has undergone extensive evolutionary processes (Foster and Kokko, 2006; Rich et al., 2021), with its establishment and development intricately regulated by a set of symbiotic genes and signaling molecules influenced by both biotic and abiotic factors (Maclean et al., 2017; Shi et al., 2021). For instance, in AM plants, the two pathways for P uptake—the direct P pathway via root hairs and epidermis and the mycorrhizal P pathway via AMF hyphae—are coordinately regulated by the same phosphate-sensing pathway (Shi et al., 2021; Das et al., 2022). Under P-deficiency conditions, host plants synthesize and release strigolactones into the soil, activating the mycorrhizal P pathway by stimulating the elongation and branching of AMF hyphae (Mori et al., 2016; Yu et al., 2022). Additionally, the extent of root colonization and the benefits conferred by AMF largely depend on the host plant species and even their genotypes (Watts-Williams et al., 2019; Huang et al., 2020). Some species are classified as “responsive plants”, which exhibit positive growth responses across most genotypes (e.g., alfalfa and maize), while others are referred to “nonresponsive plants”, exhibiting minimal or no growth enhancement but showing increased AM-activated defense responses in many genotypes (e.g., rice and wheat) (Hata et al., 2010; Li et al., 2024).

AMF are considered obligate symbionts that rely on living host plants to complete their life cycle by producing daughter spores (Smith and Read, 2008), which presents challenges for basic research on AM and their applications in agriculture and other fields. Despite recent progresses (Tanaka et al., 2022), achieving a completely asymbiotic culture of AMF—specifically, a host-free culture of daughter spores that retains the same physiological and genetic traits as their mother spore—has not been successful, partially due to their dependence on plant-derived symbiotic fatty acids (Bravo et al., 2017; Luginbuehl et al., 2017; Kameoka et al., 2019). Genomic and transcriptomic analyses have shown that AMF lack the genes required for the biosynthesis of long-chain fatty acids, and subsequent studies have demonstrated that fatty acids are transferred from host plants to AMF as their primary C source (Bravo et al., 2017; Jiang et al., 2017; Keymer et al., 2017; Luginbuehl et al., 2017). However, the mechanism underlying the allocation of plant-derived fatty acids to fungi in AM symbiosis remains unclear. In host plants, AMF colonization triggers the expression of lipid biosynthesis and transport genes (Bravo et al., 2017), with lipid transport to the AM symbiotic interface relying on the periarbuscular membrane-localized transporters stunted arbuscule (STR) and STR2 (Jiang et al., 2017; Keymer et al., 2017), and fungal uptake of plant-derived lipids mediated by fatty acid transporter 1 (RiFAT1) and RiFAT2 in the model AMF species *Rhizophagus irregularis* (Brands and Dörmann, 2022). This symbiotic C allocation is functionally linked to fungal P delivery within a framework of mutual nutritional rewards (Kiers et al., 2011), which have been disrupted during AMF-nonhost interactions due to the strong AM-activated defense responses in non-host roots (Wang et al., 2023).

Fatty acids and their derivatives, essential structural components and energy sources for plant, animal, and microbial cells, are commonly found in soil, particularly in the rhizosphere, where plant-derived fatty acids are made available through root exudates and rhizodeposition (Swenson et al., 2018; Dai et al., 2022; Liu et al., 2024). Following the discovery that AMF acquire fatty acids from their hosts during symbiosis, the use of fatty acids in cultivating asymbiotic AMF has attracted considerable attention. Kameoka et al. (2019) demonstrated that under asymbiotic conditions, exogenous palmitoleic acid and several other fatty acids stimulated the hyphal elongation and branching in *Rhisophagus irregularis* and *R. clarus,* but not in *Gigaspora margarita*, resulting in the formation of infection-competent but morphologically and functionally incomplete spores with reduced size, lower germination rates, and decreased infection capacity compared to symbiotic spores. Additionally, Sugiura et al. (2020) showed that exogenous myristate (tetradecanoic acid)—a common organic acid in various plant root exudates (Li et al., 2017; Liu et al., 2024) and soil substrates (Swenson et al., 2018; Dai et al., 2022)—can serve as a C source for the asymbiotic growth of *R. irregularis* and *R. clarus,* promoting hyphal growth, the formation of branched absorbing structures, and the production of morphologically incomplete, infection-competent spores. Subsequent studies revealed that combining plant hormones with myristate enhanced the efficiency of myristate as a C source in asymbiotic *R. clarus*, leading to the generation of numerous infection-competent spores that were smaller than those generated under symbiotic conditions (Sugiura et al., 2020; Tanaka et al., 2022). These discoveries have significantly advanced the pure culture of AMF and, importantly, identified myristate as the first external C source accessible to AMF under asymbiotic conditions, providing exceptional opportunities for understanding and utilizing the ecological functions of AM symbiosis in plant community regulation and global C cycling (Rillig et al., 2020).

However, the effects of external myristate on the growth and development of symbiotic AMF—the typical state of functioning AMF in nature—remain unclear. Regarding sugars, another type of host-derived C source, while evidence suggests that AMF extraradical hyphae in symbiosis with a carrot hairy root may absorb exogenous monosaccharides (Helber et al., 2011), the impact of such uptake on the development of symbiotic AMF remains uncertain. Moreover, no nutritional roles were observed in supporting asymbiotic spore growth following the exogenous application of sugars (Sugiura et al., 2020). A recent study reported that while exogenous myristate does not affect the extraradical biomass of symbiotic *R. irregularis* and *R. intraradices*, it enhances their colonization rates by stimulating the growth and development of extraradical hyphae. Based on this, the authors inferred that symbiotic AMF may acquire exogenous myristate (Liu et al., 2023). To date, the effects of exogenous myristate on AM symbiosis and their underlying mechanisms remain limited. Specifically, whether symbiotic AMF can uptake exogenous myristate via extraradical hyphae in the presence of plant-derived symbiotic C is yet to be determined. This question is crucial for understanding the symbiotic mechanisms and ecological roles of AM symbiosis (Rillig et al., 2020).

To address this significant knowledge gap, this study investigated the ability of symbiotic AMF to absorb external non-symbiotic myristate and examined the effect of exogenous myristate on symbiotic C-P exchanges in AM. Specifically, we aimed to determine: (i) whether symbiotic AMF can take up exogenous non-symbiotic myristate, (ii) how this additional non-symbiotic C source influences the symbiotic C allocation from plants to AMF, and (iii) if and how fungal P transfer to the host plant is affected by exogenous myristate. This work provides, to our knowledge, the strongest evidence to date that symbiotic AMF can absorb non-symbiotic fatty acids like myristate in the presence of plant-derived symbiotic carbohydrates and lipids. Furthermore, our findings reveal that exogenous myristate may, unexpectedly, disrupt the C-P exchange in AM symbiosis.

## RESULTS

### Tracing of ^13^C in AMF extraradical hyphae (Exp. 1)

To investigate the direct uptake of exogenous myristate by symbiotic AMF, we conducted an isotopic tracing experiment using ¹³C_1_-labeled myristate within the two-compartment AMF-carrot hairy root co-cultivation system (Exp. 1, Fig. 1a). Exogenous myristate specifically induced the formation of branched absorbing structures in *R. irregularis*, *R. intraradices*, and *R. diaphanus* in the hyphal compartment (HC) approximately two weeks after Myr^+^ treatment (Fig. 1b). For *R. irregularis*, the ^13^C/^12^C ratio of extraradical hyphae in HC (trial 1: 1.1504 ± 0.0575%, trial 2: 1.0919 ± 0.0001%) and root compartment (RC, trial 1: 1.1026 ± 0.0055%, trial 2: 1.0918 ± 0.0002%) under ^13^C-Myr treatment was slightly but significantly higher than that in the corresponding HC (trial 1: 1.0922 ± 0.0003%, trial 2: 1.0908 ± 0.0003%) and RC (trial 1: 1.0923 ± 0.0002%, trial 2: 1.0908 ± 0.0002%) from the non-labeled control (^12^C-Myr), respectively (*P* < 0.05, Fig. 1c). Similarly, for *R. intraradices* and *R. diaphanus*, the ^13^C/^12^C ratio of extraradical hyphae in HC (*R. intraradices*: 1.0933 ± 0.0002%, *R. diaphanus*: 1.0922 ± 0.0005%) and RC (*R. intraradices*: 1.0927 ± 0.0005%, *R. diaphanus*: 1.0930 ± 0.0004%) under ^13^C-Myr treatment was slightly but significantly higher than those in HC (*R. intraradices*: 1.0917 ± 0.0001%, *R. diaphanus*: 1.0908 ± 0.00003%) and RC (*R. intraradices*: 1.0918 ± 0.00004%, *R. diaphanus*: 1.0908 ± 0.00003%) from the non-labeled control (*P* < 0.01, Fig. 1c). Additionally, the calculated ^13^C enrichment in extraradical hyphae between HC and RC showed no significant difference for *R. irregularis* and *R. diaphanus* (Fig. 1d, e), although it was higher in HC than in RC for *R. intraradices* (*P* < 0.05), excluding the possibility that the ^13^C enrichment in AMF extraradical hyphae originated from ^13^C-myristate retention on the hyphal surface from the HC. Collectively, these results suggest that these three AMF species can take up a small amount of exogenous myristate through their extraradical hyphae when associated with host roots.

**Fig. 1.**
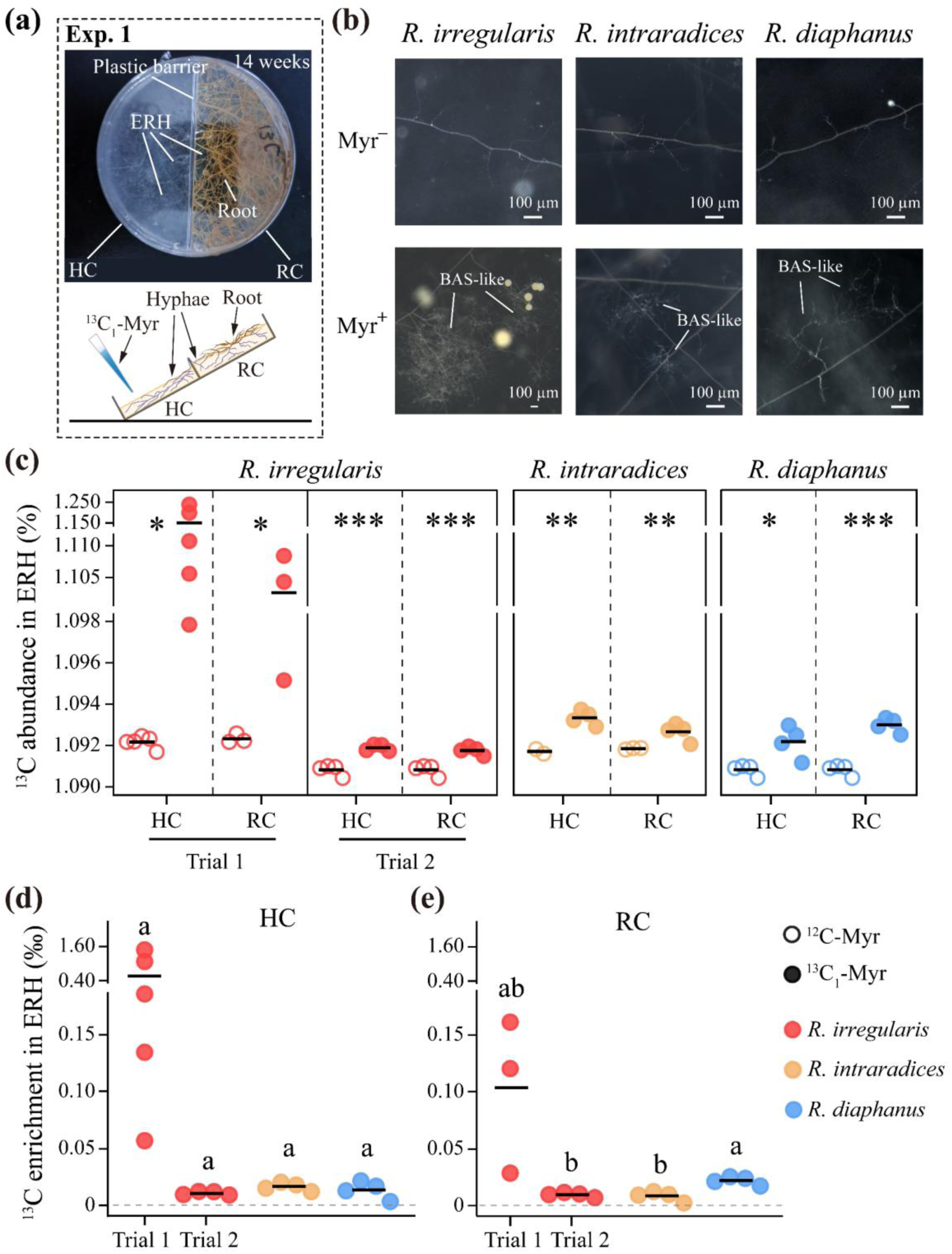
Uptake of exogenous myristate by extraradical hyphae (ERH) of arbuscular mycorrhizal fungi (AMF) under symbiotic conditions in Exp. 1. **(a)** Schematic illustration of the two-compartment AMF‒carrot hairy root co-cultivation system and the addition of myristate to the hyphae compartment (HC). Carrot hairy roots colonized by *Rhizophagus irregularis*, *R. intraradices*, or *R. diaphanus* were grown in the root compartment (RC). Twelve to fourteen weeks later, 0.1 mM of ^13^C_1_-labeled (^13^C_1_-Myr), non-labeled (^12^C-Myr) myristate, or ddH_2_O (Myr^-^, control) was added weekly to the HC with well-developed AMF extraradical hyphae (ERH) until harvest. **(b)** Branched absorbing structures (BAS) of *R. irregularis*, *R. intraradices*, and *R. diaphanus* were observed in the HC only when treated with exogenous myristate (Myr^+^). **(c)** ^13^C/^12^C ratio of the harvested ERH from HC or RC of *R. irregularis*, *R. intraradices*, and *R. diaphanus* when ^13^C_1_-labeled (^13^C_1_-Myr) or non-labeled (^12^C-Myr, background) myristate was added to HC. Two independent experimental trials were performed for *R. irregularis.* **(d, e)** Calculated ^13^C enrichment in ERH of *R. irregularis*, *R. intraradices*, and *R. diaphanus* in the HC and RC under ^13^C-Myr treatment. Black horizontal lines represent mean value from 2‒5 biological replicates. *, **, and *** indicate statistical significance between ^13^C_1_-labeled and non-labeled treatments at the 0.05, 0.01, and 0.001 probability levels (Mann-Whitney U-test), respectively. Different letters indicate significant difference at the 0.05 probability level (one-way ANOVA followed by Games-Howell’s post hoc test). Source data are provided as a Source Data file.

The ¹³C enrichment levels among the three AMF species were calculated to compare their capacity for myristate absorption. In HC, the calculated ^13^C enrichments in extraradical hyphae of *R. irregularis* (trial 1: 0.5823 ± 0.4111‰, trial 2: 0.0107 ± 0.0014‰), *R. intraradices* (0.0163 ± 0.0031‰), and *R. diaphanus* (0.0136 ± 0.0067‰) showed no significant differences in the ^13^C-Myr treatment (Fig. 1d). In RC, the ^13^C enrichment in extraradical hyphae also showed no significant differences (Fig. 1e), except for a slightly but significantly higher ^13^C enrichment in *R. diaphanus* (0.0217 ± 0.0031‰) compared to *R. intraradices* (0.0082 ± 0.0037‰) and *R. irregularis* in trial 2 (0.0093 ± 0.0017‰, *P* < 0.05). These results suggest a generally similar myristate absorption capacity among these AMF species under the tested conditions.

### Effects of myristate on root colonization and expression of fatty acid uptake and metabolism marker genes (Exp. 1)

The expression of fatty acid uptake and metabolism marker genes in *R. irregularis* extraradical hyphae were analyzed to investigate their uptake of myristate from HC at the transcriptional level. The results showed that exogenous myristate significantly increased the expression levels of the fungal fatty acid transporter gene *RiFAT2* (*P* < 0.05), while had no significant effect on *RiFAT1* (Fig. 2a). Additionally, exogenous myristate significantly or marginally significantly increased the expression levels of the fungal genes involved in fatty acid β-oxidation (*RiFAD1*, *P* < 0.05), palmitvaccenic acid synthesis (*RiOLE1*, *P* = 0.057), and TCA cycle activity (*RiCIT1*, *P* < 0.05) (Fig. 2b). These results suggest that exogenous myristate activates the expression of fungal genes related to fatty acid uptake and metabolism in the extraradical hyphae of *R. irregularis*. For AMF colonization intensity, exogenous myristate significantly increased the hyphal and arbuscular colonization rates of *R. irregularis, R. intraradice*, and *R. diaphanus* in carrot hairy roots (*P* < 0.05, Fig. 2c−e) but had no effect on vesicle colonization.

**Fig. 2.**
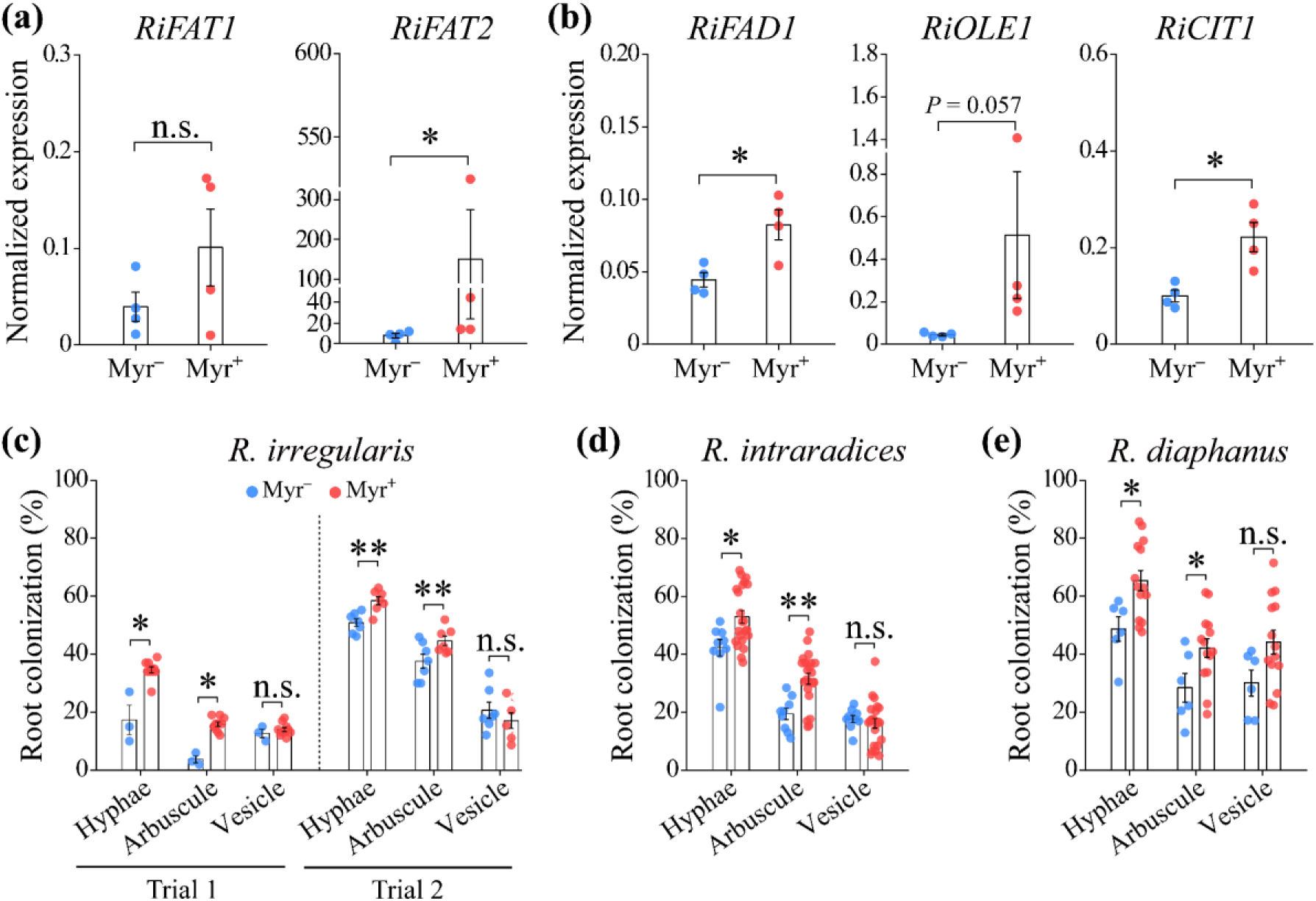
Effects of exogenous myristate on the expression of fungal fatty acid uptake and metabolism genes in the extraradical hyphae (ERH) and on root colonization by arbuscular mycorrhizal fungi (AMF) in Exp. 1. (**a, b**) Normalized expression (relative to *RiEF-1β*) of AMF genes involved in **(a)** fatty acid uptake (*RiFAT1* and *RiFAT2*) and **(b)** metabolism—β-oxidation (*RiFAD1*), palmitvaccenic acid synthesis (*RiOLE1*), and TCA cycle (*RiCIT1*)—in the ERH of *R. irregularis* collected from the hyphae compartment (HC) under myristate (Myr^+^) or ddH_2_O (Myr^-^) treatments (n = 4). **(c−e)** Percent root colonization by **(c)** *R. irregularis* (n = 3‒10), **(d)** *R. intraradices* (n = 9‒22), and **(e)** *R. diaphanus* (n = 6‒14) in carrot hairy roots from the root compartment (RC) when HC was treated with myristate (Myr^+^) or ddH_2_O (Myr^-^). Two independent experimental trials were performed for *R. irregularis*. Means ± SE, *, **, and *** indicate statistical significance between Myr^+^ and Myr^-^ treatments at the 0.05, 0.01, and 0.001 probability levels, respectively. Source data are provided as a Source Data file.

### Effects of exogenous myristate on AMF spore germination, root colonization, and extraradical structure development (Exp. 2)

The impact of exogenous myristate on the growth and development of symbiotic AMF was evaluated in a single-compartment *R. irregularis*-carrot hairy root co-cultivation system (Exp. 2, Fig. 3a). Exogenous myristate significantly promoted spore germination (*P* < 0.001, Fig. 3b), and enhanced hyphal branching and length (*P* < 0.001, Fig. 3c, d) before visible physical contact between hyphae and hairy roots at four days after AMF inoculation. Morphological examinations showed that exogenous myristate significantly enhanced the colonization intensities of hyphae and arbuscules, but not vesicles, in carrot hairy roots across various application levels (Fig. 3e). Exogenous myristate also significantly increased the extraradical biomass of AMF (hyphae and spore) at different application concentrations (*P* < 0.05, Fig. 3f).

**Fig. 3.**
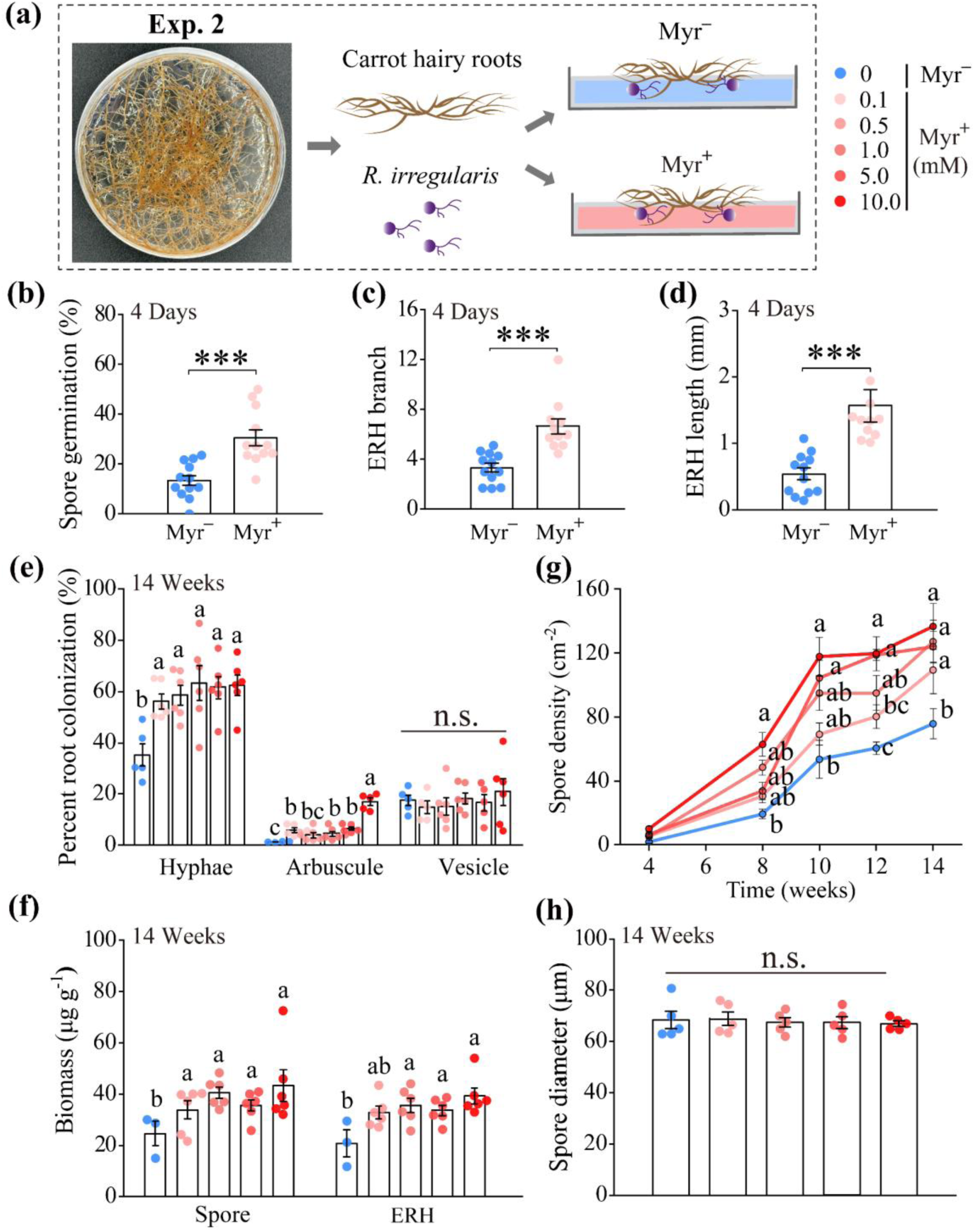
Effects of exogenous myristate on the development of intraradical and extraradical structures of AMF (*Rhizophagus irregularis*) in the single-compartment AMF-carrot hairy root co-cultivation system (Exp. 2). **(a)** Schematic illustration of the single-compartment *R. irregularis*-hairy root co-cultivation system and the addition (Myr^+^) or absence (Myr^-^) of myristate. **(b)** Spore germination rate, **(c)** extraradical hyphae (ERH) branching, and **(d)** ERH length in the *R. irregularis*-hairy root co-cultivation system with (Myr^+^) or without (Myr^-^) myristate treatment at 4 days post-inoculation (n = 11−12). **(e)** AMF percent root colonization in carrot hairy roots and **(f)** biomass of the AMF extraradical spore and hypha (per gram of medium) at 14 weeks post-inoculation (n = 3−6). **(g)** Dynamics of AMF spore density over time (per square centimeter, n = 3−5); and **(h)** AMF spore diameter at harvest with 0, 0.1, 0.5, 1, 5, or 10 mM myristate treatment under symbiotic conditions (n = 5, calculation from > 900 spores in each replicate). Means ± SE; different letters indicate significant differences at *P* < 0.05 level (one-way ANOVA followed by Tukey’s post hoc test); *** indicates statistical significance at the 0.001 probability level (Student’s *t*-test). Source data are provided as a Source Data file.

Meanwhile, exogenous myristate significantly increased the density of AMF spores from eight weeks post-AMF inoculation (*P* < 0.05, Fig. 3g), and the diameter of newly produced AMF spores in media with exogenous myristate was identical to those in media without myristate (Fig. 3h).

### Environmental myristate levels and the effects of exogenous myristate on AMF intraradical and extraradical biomass during symbiosis with alfalfa and rice (Exp. 3)

In Exp. 3 (Fig. 4a), our field survey on the environmental myristate distributions revealed that myristate was extensively detected in soil and plant samples from the paddy field, grassland, and woodland at a 2.87‒14.68 mg kg^-1^ level (Fig. 4b). On average, the myristate level in plant (fresh leaf or root) is mostly higher than that in soil (leaf litter, rhizosphere soil, or bulk soil) (*P* < 0.05, Fig. 4b). Additionally, the myristate levels in leaf litter and rhizosphere soil were usually greater than in bulk soil (*P* < 0.05), suggesting that plants are significant source of myristate in the soil. The myristate contents from the same sample type typically varied across different habitats (*P* < 0.05). A pot experiment was further conducted to assess the effect of exogenous myristate on C-P nutritional exchanges in AM symbiosis (Exp. 3). All typical AMF intraradical structures (hyphae, arbuscules, and vesicles) were observed in the roots of both alfalfa and rice in the *R. irregularis*-inoculated groups (AM), while no AMF colonization was detected in the non-inoculated (NM) groups. Under low-P conditions, exogenous myristate significantly increased arbuscule abundance in alfalfa roots (*P* < 0.05, Fig. 4c) but did not significantly affect hyphae or vesicle abundance (Supporting Information Fig. S1a, c). Exogenous myristate significantly enhanced arbuscule, hyphae, and vesicle abundance in rice roots (*P* < 0.05, Fig. 4e, Supporting Information Fig. S1b, d). However, under high-P conditions, exogenous myristate had no effect on AMF abundance in either alfalfa or rice roots (Fig. 4, Supporting Information Fig. S1). AMF signature fatty acid (NLFA 16:1ω5) and the relative expression of the AMF marker gene elongation factor-1 beta (*RiEF-1β*) were analyzed as indicators of AMF biomass. For intraradical AMF biomass, exogenous myristate significantly increased the levels of the NLFA 16:1ω5 and the *RiEF-1β* expression in both alfalfa and rice roots under low-P conditions (*P* < 0.05, Fig. 4d, f, g, h); however, it had no significant impact under high-P conditions. In alfalfa roots, NLFA 16:1ω5 levels were higher under high-P than low-P conditions (*P* < 0.05). For extraradical AMF biomass, exogenous myristate increased NLFA 16:1ω5 levels in the rhizosphere soils of alfalfa (*P* < 0.05) but not in rice (*P* = 0.14) under low-P conditions (Fig. 4g, h). Under high-P conditions, myristate had no effect on NLFA 16:1ω5 levels in the rhizosphere soil of either plant.

**Fig. 4.**
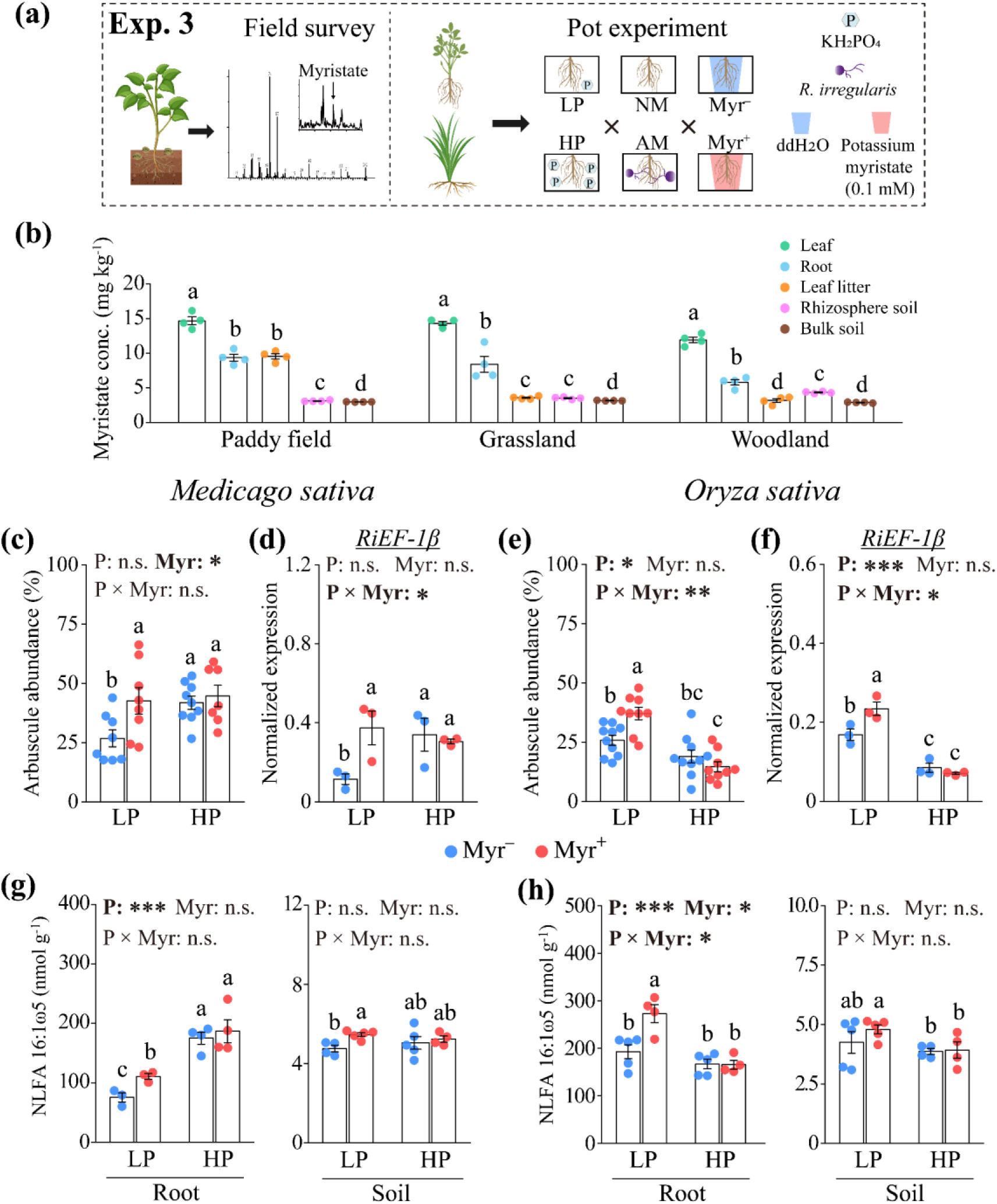
Environmental myristate levels and the effects of exogenous myristate on arbuscule abundance, as well as the intraradical and extraradical biomass of arbuscular mycorrhizal fungi (AMF, *Rhizophagus irregularis*) associated with *Medicago sativa* or *Oryza sativa* under low (LP) or high (HP) phosphorus conditions (Exp. 3). **(a)** Schematic illustration of the field survey of environmental myristate levels and experimental design in the pot experiment (Exp. 3). **(b)** Myristate concentration in bulk soil, as well as in plant leaves, roots, leaf litter, and rhizosphere soil from the dominant species *O. sativa*, *Axonopus compressus*, and *Morus alba* in the paddy field, grassland, and woodland, respectively (n = 4). **(c, e)** Arbuscule abundance in *M. sativa* and *O. sativa* roots treated with 0.1 mM myristate (Myr^+^) or ddH_2_O (Myr^-^) (n = 10; vesicle and hyphal abundance are shown in Supporting Information Fig. S1); **(d, f)** Normalized expression of *RiEF-1β* (relative to *MsACT2* in *M. sativa* and *OsCyc2* in *O. sativa*) in AMF-colonized roots with 0.1 mM myristate (Myr^+^) or ddH_2_O (Myr^-^) treatments (n = 3). **(g, h)** AMF biomass, indicated by the AMF signature fatty acid (NLFA 16:1ω5), in AMF-colonized roots and rhizosphere soil of *M. sativa* and *O. sativa* treated with myristate (Myr^+^) or ddH_2_O (Myr^-^) (n = 5). Means ± SE; different letters indicate significant differences at *P* < 0.05 level (**b**: Student’s *t*-test, **c**−**h**: two-way ANOVA followed by Tukey’s post hoc test). Source data are provided as a Source Data file.

### Effects of exogenous myristate on the transfer of symbiotic C to AMF (Exp. 3)

The analysis of ^13^C-NLFA-isotope ratio mass spectrometry was performed to investigate the impact of exogenous myristate on the allocation of symbiotic C from plants to AMF. In AM plants, there was a notable enrichment of ^13^C (δ^13^C = 1.75 ± 0.6‰) in the AMF signature fatty acid (NLFA 16:1ω5) compared to NM plants (δ^13^C = −24.8 ± 2.3‰), demonstrating the transfer of newly fixed ^13^C from plants to AMF. The calculated ^13^C flow from alfalfa to root-colonizing AMF (0.3400‒1.2642 μg) was over ten times higher than that from rice (0.0261‒0.0596 μg), highlighting a substantially greater efficiency of plant-derived symbiotic C delivery to AMF in AM-responsive alfalfa compared to non-responsive rice. In alfalfa, exogenous myristate reduced the symbiotic C transfer to AMF (Myr effect: *P* < 0.001, Fig. 5a), although this effect was not significant under high-P conditions. In rice, myristate also significantly influenced the symbiotic C flow to AMF, with its effect significantly interacting with soil P level (Myr × P: *P* = 0.004, Fig. 5b): myristate significantly increased the allocation of ^13^C to AMF under low-P conditions (*P* < 0.05) but decreased it under high-P conditions (*P* < 0.01).

**Fig. 5.**
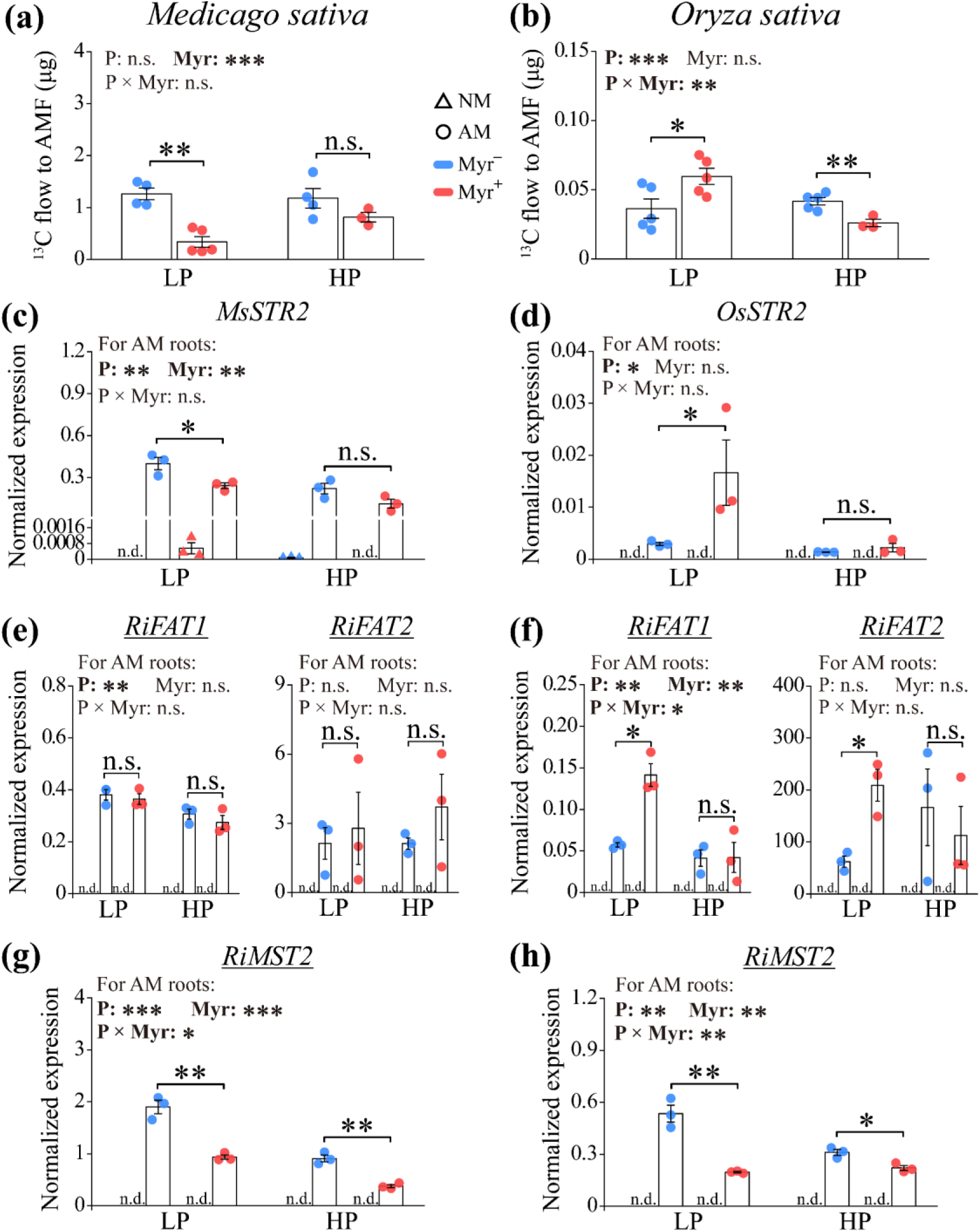
Effects of exogenous myristate on symbiotic carbon (C) allocation from *Medicago sativa* and *Oryza sativa* to arbuscular mycorrhizal fungi (AMF, *Rhizophagus irregularis*) and on the expression profiles of mycorrhizal C transportation marker genes. **(a, b)** Plant-derived ^13^C flow to the AMF signature fatty acid (NLFA 16:1ω5) in AMF-colonized roots of *M. sativa* and *O. sativa* treated with 0.1 mM myristate (Myr^+^) or ddH_2_O (Myr^-^) under low (LP) and high (HP) phosphorus conditions (n = 5, two-way ANOVA tested the effects of the treatments and their interaction). (**c**−**h**) Normalized expression of plant and fungal (underlined) marker genes for mycorrhizal C transport in AMF-colonized (AM) and non-colonized (NM) roots treated with 0.1 mM myristate (Myr^+^) or ddH_2_O (Myr^-^) (n = 3). The mycorrhizal C transportation genes include the plant symbiotic fatty acid transporter *STR2* **(c, d**), the fungal fatty acid transporters (*RiFAT1, RiFAT2*; **e, f**), and the monosaccharide transporter (*RiMST2*; **g, h**). Normalized expression of *M. sativa*, *O. sativa*, and AMF genes was calculated relative to *MsACT2*, *OsCyc2*, and *RiEF-1β*, respectively; n.d., not detected. In c−h, the two-way ANOVA in each histogram applies only to AM roots. Means ± SE; * and ** indicate statistical significance at the 0.05 and 0.01 probability levels, respectively (Student’s *t*-test). Source data are provided as a Source Data file.

At the transcriptional level, the qRT-PCR analysis revealed that the expressions of the symbiotic lipid transfer marker genes *STR2* (in alfalfa and rice) and the fungal genes responsible for symbiotic fatty acid uptake *(RiFAT1* and *RiFAT2)* or monosaccharide uptake (*Monosaccharide Transporter 2* (*RiMST2*)) were exclusively detected in AM roots, except for a low expression level of alfalfa *MsSTR2* occasionally observed in NM roots (Fig. 5c–h). In AMF-inoculated alfalfa roots, exogenous myristate significantly downregulated the expressions of the symbiotic C transfer genes *MsSTR2* in the plant and *RiMST2* in AMF (*P* < 0.05), while having no effect on *RiFAT1* and *RiFAT2* in AMF (Fig. 5c, g, e). In AMF-inoculated rice roots, the effects of myristate on the tested symbiotic C transfer genes were largely influenced by soil P levels (*P* < 0.05). Under low-P conditions, myristate significantly increased the expressions of plant *OsSTR2* and fungal *RiFAT1* and *RiFAT2*, while decreasing the expression of fungal *RiMST2* (*P* < 0.05, Fig. 5d, f, h). Under high-P conditions, myristate significantly increased the expression of *RiMST2* (*P* < 0.05) but had no effects on *OsSTR2*, *RiFAT1*, and *RiFAT2*. In both plant species with AMF inoculation, soil P level showed significant effects on *STR*, *RiFAT1*, and *MST2* genes (*P* < 0.05), with generally lower expression levels of these genes under high-P compared to low-P conditions.

### Effects of exogenous myristate on the mycorrhizal P pathway and mycorrhizal growth response (Exp. 3)

We further analyzed the mycorrhizal P pathway and mycorrhizal growth response in the pot experiment to evaluate the effects of exogenous myristate on AM symbiosis (Exp. 3). AMF inoculation significantly increased P concentrations in both alfalfa and rice plants (*P* < 0.001, Fig. 6a, b). Myristate generally exhibited no effects on plant P concentrations for AM and NM plants, except that it significantly decreased P concentration in AMF-colonized alfalfa under high-P conditions (*P* < 0.05). Under low-P conditions, exogenous myristate significantly decreased the mycorrhizal P response in both alfalfa and rice (*P* < 0.05, Fig. 6c, d). Under high-P conditions, while exogenous myristate significantly decreased the mycorrhizal P response in alfalfa (*P* < 0.05), it had no significant effect on rice.

**Fig. 6.**
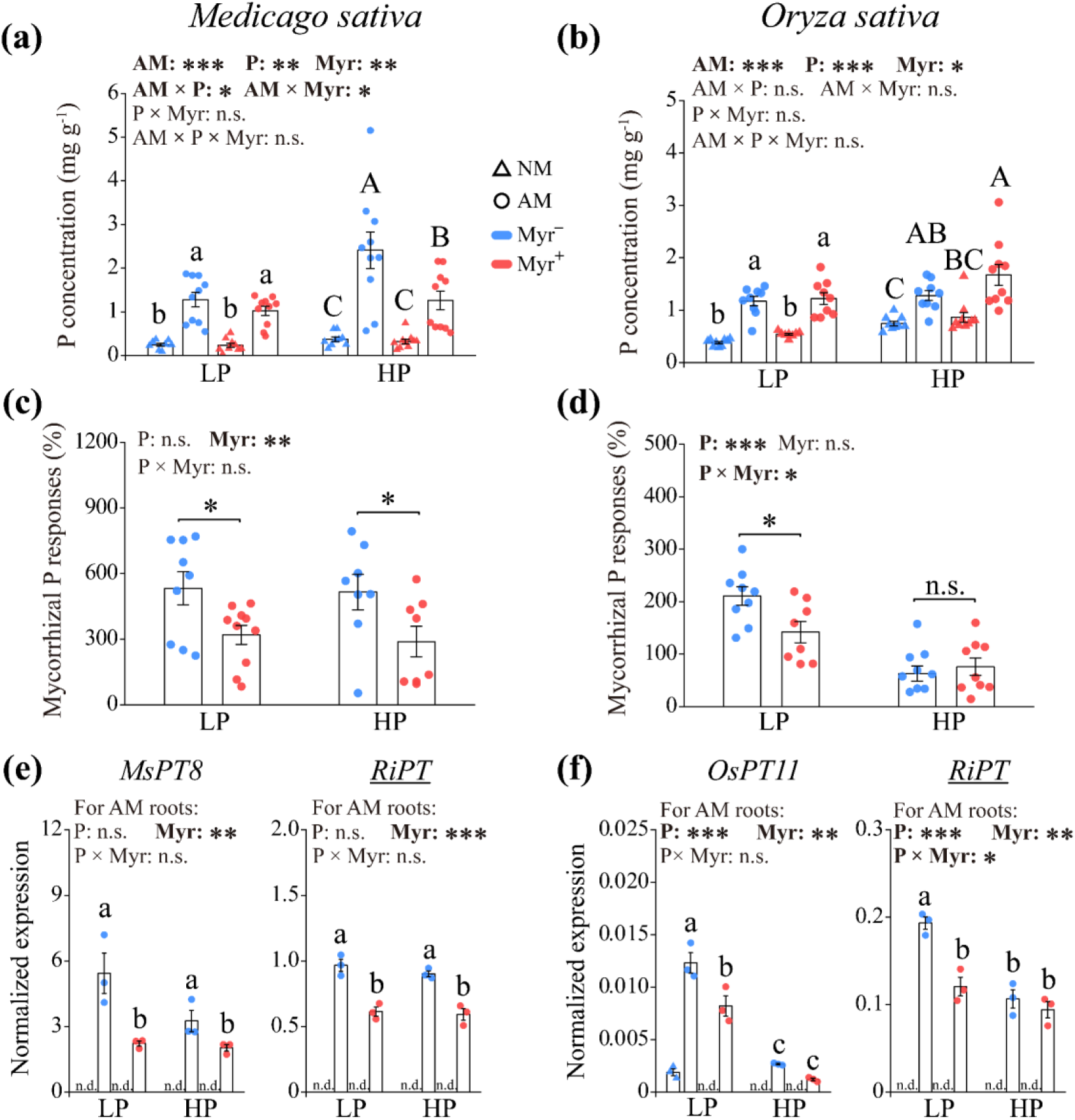
Effects of exogenous myristate on plant phosphorus (P) concentration, mycorrhizal P response, and expression of plant and fungal (underlined) marker genes for the mycorrhizal P pathway in *Medicago sativa* and *Oryza sativa*. **(a, b)** Plant P concentration (n = 8−10); **(c, d)** mycorrhizal P response (n = 8−10); and **(e, f)** relative expression of *MsPT8*, *OsPT11*, and *RiPT* in *M. sativa* and *O. sativa* roots under low (LP) and high (HP) phosphorus conditions with 0.1 mM myristate (Myr^+^) or ddH_2_O (Myr^-^) treatments (n = 3). AM, inoculated with *Rhizophagus irregularis*; NM, non-inoculated; n.d., not detected. Normalized expression of *M. sativa*, *O. sativa*, and AMF genes was calculated relative to *MsACT2*, *OsCyc2*, and *RiEF-1β*, respectively. In a−d, two-or three-way ANOVA tested the effects of the treatments and their interaction; In e and f, two-way ANOVA in each histogram applies only to AM roots, followed by Tukey’s post hoc test for multiple comparisons. Means ± SE; different letters indicate significant differences at *P* < 0.05 level. * and *** indicate statistical significance at the 0.05 and 0.001 probability levels (Student’s *t*-test), respectively. Source data are provided as a Source Data file.

At the transcriptional level, the expression of mycorrhizal P pathway marker genes in plants (*Phosphate Transporter*, *MsPT8* or *OsPT11*) and AMF (*RiPT*) was detected exclusively in the AM roots of rice and alfalfa, except for a low expression level of rice *OsPT11* observed in the NM root under low-P and Myr^−^ conditions. This finding confirmed that AM symbiosis activated the mycorrhizal P pathway at the transcriptional level in both plant species (Fig. 6e, f). Exogenous myristate significantly influenced the transcriptional responses of the mycorrhizal P pathway in both alfalfa and rice (*P* < 0.01). In alfalfa, exogenous myristate downregulated the expressions of *MsPT8* and *RiPT* under both low-and high-P conditions (*P* < 0.05, Fig. 6e). In rice, exogenous myristate decreased the expression levels of *OsPT11* and *RiPT* under low-P conditions (*P* < 0.05), but not under high-P conditions (Fig. 6f).

In alfalfa, AMF inoculation significantly increased plant biomass under both low-and high-P conditions (*P* < 0.05, Supporting Information Fig. S2a), with mycorrhizal growth responses significantly higher under high-P than low-P conditions (*P* < 0.05, Supporting Information Fig. S2c). In rice, AMF inoculation had no effect on plant biomass under low-P conditions, but reduced rice biomass under high-P conditions (*P* < 0.05); mycorrhizal growth responses were significantly lower under high-P than low-P conditions (*P* < 0.05, Supporting Information Fig. S2b, d). Exogenous myristate had no effect on plant biomass or mycorrhizal growth response in either alfalfa or rice, while soil P levels significantly influenced mycorrhizal growth response: high-P treatment increased mycorrhizal growth response in alfalfa but decreased it in rice compared to low-P conditions (*P* < 0.01, Supporting Information Fig. S2c, d).

### Effects of exogenous myristate on the transcriptome of AM roots (Exp. 3)

Transcriptome analysis was performed to preliminarily explore the effects of exogenous myristate on mycorrhizal roots at the transcriptional level. The RNA-seq data identified 39,944 and 33,056 genes (transcripts) in alfalfa and rice, respectively. Principal component analysis of gene expression profiles in mycorrhizal roots of alfalfa and rice showed that the first and second principal components together explained 32.4% and 44.1% of the variation, respectively (Supporting Information Fig. S3a, b). In both alfalfa and rice roots, gene expression profiles with or without myristate application could be partially separated under low-P conditions, but not under high-P conditions, suggesting the importance of P status in myristate’s effect.

Exogenous myristate induced a variety of differentially expressed genes (DEGs, Myr^+^ *vs.* Myr^−^) in mycorrhizal alfalfa (LP: 308, HP: 137) and rice (LP: 590, HP: 855) roots (Supporting Information Fig. S3c, d). Gene Ontology (GO) analysis of these DEGs enriched several GO terms related to plant stress or defense, most of which were downregulated by exogenous myristate (Fig. 7a, b). These included “defense response”, “response to salicylic acid”, “peroxidase activity”, etc., in rice, and “glycosyltransferase activity” and “alternative oxidase activity” in alfalfa, suggesting that myristate may modulate the normal plant defense response in AM symbiosis.

**Fig. 7.**
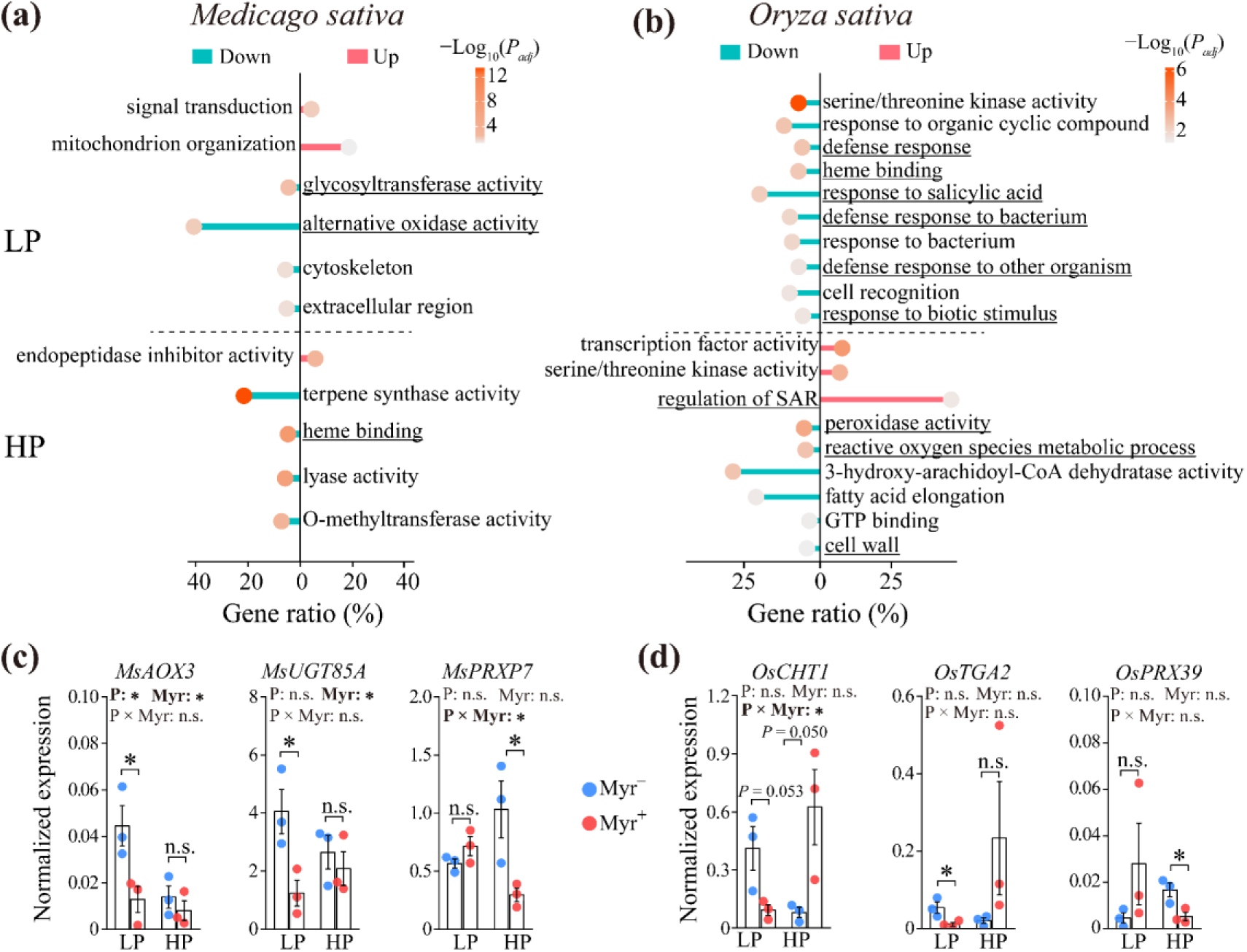
Effects of exogenous myristate on the global transcriptome and putative AM-activated defense genes in *Medicago sativa* and *Oryza sativa* roots inoculated with arbuscular mycorrhizal fungi (AMF, *Rhizophagus irregularis*) under low (LP) and high (HP) phosphorus conditions. **(a, b)** Significantly enriched gene ontology (GO) terms (*P_adj_* < 0.05) for differentially expressed genes (DEGs) between myristate (Myr^+^) and ddH_2_O (Myr^-^) treatments in RNA-seq. Underlined terms denote stress-or defense-related GO categories, with the expression profile of their component genes provided in Supporting Information (Fig. S4). **(c, d)** Quantitative real-time PCR analysis of the AM-activated defense gene expression—reactive oxygen species (ROS) balance (*MsAOX3*), stress responses (*MsUGT85A*, *MsPRXP7*, and *OsPRX39*), and defense responses *(OsCHT1* and *OsTGA2*) — between Myr^+^ and Myr^-^ treatments under LP and HP conditions (n = 3, two-way ANOVA tested the effects of the treatments and their interaction). Means ± SE; * indicates statistical significance between Myr^+^ and Myr^-^ treatments at the 0.05 probability level (Student’s *t*-test). Source data are provided as a Source Data file.

To further characterize this response, we subsequently analyzed the expression profiles of the component genes associated with enriched GO terms that are related to defense or stress, using the RNA-seq dataset (Fig. S4a, b). In alfalfa, exogenous myristate significantly downregulated several key genes involved in reactive oxygen species balance (*alternative oxidase 2* (*AOX2*), *AOX3*) and stress tolerance (*UDP-glycose flavonoid glycosyltransferase* (*UFGT*), *UDP-glycosyltransferase* (*UGT*) *73C3*, *UGT73C6*) under low-P conditions, as well as typical stress-responsive genes (*peroxidase* (*PRX*) *P7*, *cytochromes P450* (*CYP*) *83B1*), and gene families such as *geraniol 8-hydroxylase-like* (*G8HL*), *CYP71D10*, and *CYP736A12* under high-P conditions (Supplementary Fig. S4a). In rice, exogenous myristate downregulated several typical defense-associated transcriptional factors (*TGA2*, *TGA4*, *WRKY45*, *WRKY62*, *WRKY76*, and *bZIP79*) and marker genes (*pathogenesis-related 5-like* (*PR5L*), *protein phosphatase 2C 35* (*OsPP2C35*), *1-aminocyclopropane-1-carboxylate synthase 2* (*ACS2*), *DWARF4*, *pyricularia oryzae resistance-ta* (*PITA*), *mildew resistance locus O2* (*MLO2*), *NRR repressor homologue 1* (*RH1*), and *chitinase 1* (*CHT1*)) under low-P conditions (Supplementary Fig. S4b). However, most of these genes were upregulated following myristate application under high-P conditions. Collectively, these results indicate that exogenous myristate modified AM-activated plant defense response in plants, although this effect is influenced by host identity and soil P levels.

### qRT-PCR verification of the effects of exogenous myristate on AM-activated defense genes in mycorrhizal roots (Exp. 3)

To verify the impact of exogenous myristate on the AM-activated defense response, qRT-PCR was performed to analyze the expression of defense marker genes in both plant species. In alfalfa roots, exogenous myristate significantly decreased the expression of *alternative oxidase 3 (MsAOX3)* and *UDP-glycosyltransferase 85A* (*MsUGT85A*) (*P* < 0.05), while its effect on *peroxidase P7* (*MsPRXP7*) depended on soil P level (Myr × P: *P* < 0.05): exogenous myristate significantly decreased *MsPRXP7* expression under high-P conditions (*P* < 0.05) but had no effect under low-P conditions (Fig. 7c). In rice roots, the effect of exogenous myristate on *chitinase 1* (*OsCHT1*) was also dependent on soil P level (Myr × P: *P* < 0.05): it marginally decreased *OsCHT1* expression under low-P conditions (*P* = 0.053) but increased it under high-P conditions (*P* = 0.050). Exogenous myristate significantly suppressed the expression of defense-related transcriptional factor *OsTGA2* and *peroxidase 39* (*OsPRX39*) under low-P and high-P conditions, respectively (*P* < 0.05, Fig. 7d). Overall, the qRT-PCR results largely confirmed the inhibitory effects of myristate on AM-activated defense responses in alfalfa and low-P rice.

## DISCUSSION

AMF are obligate biotrophs that rely on host-derived symbiotic lipids for reproduction (Bravo et al., 2017; Jiang et al., 2017; Keymer et al., 2017; Luginbuehl et al., 2017). However, whether symbiotic AMF can uptake external non-symbiotic lipids in the presence of plant-derived C remains poorly understood (Rillig et al., 2020; Liu et al., 2023). Under asymbiotic conditions, Sugiura et al. (2020) found that among eight saturated or unsaturated fatty acids (C12 to C18) and two β-monoacylglycerols supplied, only myristate led to an increase in the biomass of *R. irregularis*. In our study, we observed slight yet statistically significant ^13^C enrichments in the extraradical hyphae of three AMF species from both hyphal and root compartments during the ^13^C_1_-myristate tracing experiment (Fig. 1c), indicating that the extraradical hyphae of the tested AMF species can absorb external non-symbiotic myristate in the presence of plant-derived symbiotic carbohydrates and lipids. Several additional observations further supported this finding. First, exogenous myristate specifically induced the formation of branched absorbing structures in AMF extraradical hyphae (Fig. 1b), which may serve as preferential sites for myristate uptake (Bago et al., 1998; Sugiura et al., 2020). Second, exogenous myristate activated the transcription of marker genes associated with fatty acid uptake and metabolism in extraradical hyphae (Fig. 2). Third, myristate application frequently enhanced both intraradical and extraradical AMF biomass (Figs. 2, 3, and 4), even when the symbiotic C supply from the plant was reduced (Figs. 4 and 5), accompanied by decreased mycorrhizal P responses (Fig. 6). In a priority study, myristate application also increased colonization intensity, but not the extraradical biomass of AMF in tomato roots (Liu et al., 2023). Overall, our findings indicate that the extraradical hyphae of symbiotic AMF can uptake external non-symbiotic fatty acids like myristate beyond those directly delivered by the host, at least under the conditions studied.

Notably, myristate has been frequently detected in root exudates and soil substrates from non-targeted metabolome analyses (Li et al., 2017; Swenson et al., 2018; Dai et al., 2022; Liu et al., 2024), although its specific levels in various soil environments remain largely unknown. Our field survey demonstrates that myristate is widely present in soil, litter, and plant (Fig. 4b), with its environmental level (2.87−14.68 mg kg^-1^) encompassing the myristate treatments applied in our greenhouse experiment (Exp. 3, totaling 1.83 mg per 1000 ml or 300 ml soil substrate in rice and alfalfa, respectively). Therefore, it is likely that the widespread AMF extraradical hyphae network may absorb myristate from soil environments such as root exudates, decaying plants, or litter in realistic environments. Further research is still needed to validate this hypothesis; however, it effectively explains a long-observed but poorly understood phenomenon—AMF often colonize and thrive in leaf litter and undecomposed plant material despite their limitations in assimilating organic forms of nutrients (Went and Stark, 1968; Zhang et al., 2018; Bunn et al., 2019)—as the myristate concentration in plant and leaf litter is higher than that in soil (Fig. 4b).

During AM establishment, plant roots typically activate low-level defense responses to limit excessive C consumption caused by over-colonization (Blilou et al., 2000; Feng et al., 2019). In our study, myristate increased both intraradical and extraradical biomass of the symbiotic AMF and promoted the formation of normal-sized secondary spores (Figs. 2 and 3), unlike the small spores induced by myristate under asymbiotic conditions (Sugiura et al., 2020). This suggests that plant roots supply additional substances to AMF beyond myristate, the identification of which is essential for the pure culture of AMF. Meanwhile, exogenous myristate transcriptionally downregulated plant genes associated with AM-activated defenses under low-P conditions (Fig. 7), suggesting that myristate may interfere with normal defense mechanisms in host roots to facilitate AMF colonization. This effect contrast with AM signals like strigolactones and quercetin, which enhance plant defenses (Sudheeran et al., 2020; Kusajima et al., 2022). Further research is required to clarify myristate’s role in regulating AMF colonization and development.

Despite increased arbuscule abundance (Fig. 4)—where symbiotic nutritional exchange occurs (Smith and Read, 2008)—exogenous myristate unexpectedly reduced the mycorrhizal P response and downregulated plant and fungal mycorrhizal P transporter gene expression in both alfalfa and rice roots (Fig. 6), indicating a diminished mycorrhizal P benefit through inactivation of mycorrhizal P pathway transcription. Meanwhile, exogenous myristate also reduced C supply to AMF from both alfalfa and rice, except in low-P rice (Fig. 5). Together, these results suggest that exogenous myristate disrupts the symbiotic C-P exchange, which is functionally coupled in AM symbiosis (Kiers et al., 2011). This disruption could be attributed to several factors. First, it may result from AMF partially shifting from utilizing symbiotic C to absorbing external C (myristate, Fig. 1), thus reducing their competition for symbiotic C at the AM interaction interface (Bitterlich et al., 2014). Correspondingly, this decline in symbiotic C supply to AMF may decrease mycorrhizal P benefit, since AMF can modulate P transport based on the rate at which plants supply C (Kiers et al., 2011; Argüello et al., 2016). Second, exogenous myristate may interfere with AM signaling, as implied by the interference in normal AM-activated defenses (Fig. 7), thereby limiting nutrient exchange in the symbiosis. Third, AMF P transport proteins expressed in arbuscular cells can re-uptake P from the symbiotic interface, creating P competition between the plant and AMF (Xie et al., 2022). In our study, the increased fungal biomass with exogenous myristate suggests a higher P demand by AMF, reducing the feasibility of nutritional trade.

In contrast to its effect under high-P treatment, myristate increased, rather than decreased, the allocation of rice-derived C to AMF under low-P conditions, accompanied by enhanced expression of marker genes for fatty acid (but not sugar) allocation from rice to AMF (Fig. 5b, f, g). These results highlight that myristate’s impact on AM symbiosis is influenced by soil P status, a significant regulatory factor of the symbiosis by coordinating the direct P pathway and mycorrhizal P pathway in AM plants (Shi et al., 2021; Das et al., 2022). However, the significant effects of soil P status on myristate’s impact on C-P exchange were not observed in alfalfa (Figs. 5 and 6), where the unexpectedly higher AMF intraradical biomass (Fig. 4) and greater mycorrhizal growth response (Fig. S2) in the high-P compared to the low-P treatment are likely due to the inhibitory effects of extremely low-P environments on AMF colonization and functioning (Janos, 2007; Wang et al., 2020). The varying responses of myristate’s effect to soil P treatment between rice and alfalfa may arise not only from the different status of plant P nutrition but also from their varying symbiotic strategies: AM-responsive alfalfa demonstrated significantly higher efficiency in C-P nutritional exchange compared to AM-nonresponsive rice (Figs. 5 and 6). Collectively, our findings regarding the disruptive effects of myristate on symbiotic C-P exchange, along with the broad yet varied presence of myristate in different soil components (Fig. 4), imply that myristate may be a critical influencing factor in AM symbiosis. Interestingly, myristate also exhibited inhibitory effects on rhizospheric *Devosia*, which is responsible for promoting growth and suppressing disease in *Arabidopsis* (Liu et al., 2024). Further research is required to clarify the mechanisms underlying the disruptive effect of exogenous myristate on symbiotic C-P exchange. Nevertheless, our findings advise caution against the application of myristate, and possibly other non-symbiotic C sources of AMF, to enhance AM symbiosis, as it disrupts the symbiotic nutritional equilibrium and may potentially weaken plant immunity.

Despite these encouraging findings, our work has several limitations. First, the absorption of myristate by symbiotic AMF was observed only after exogenous application under artificial conditions, which may not accurately reflect natural environments. Second, our study focused exclusively on myristate and AMF species from the genus *Rhizophagus*, although different AMF groups may respond to myristate or other fatty acids in various ways, at least under asymbiotic conditions (Kameoka et al., 2019; Sugiura et al., 2020). Third, our investigation into the mechanism by which myristate disrupts C-P exchange in AM symbiosis remains preliminary, and the limited P supply along with varied growth conditions between alfalfa and rice further complicate the mechanistic analysis. Therefore, further research is needed to verify and quantitatively assess the absorption of external myristate and other C sources by symbiotic AMF under natural conditions, as well as to elucidate their regulatory mechanisms and ecological significance in AM symbiosis.

## CONCLUSIONS

In summary (Fig. 8), our study demonstrates that symbiotic AMF can uptake external non-symbiotic fatty acids such as myristate, which is commonly present in soil components. Exogenous myristate enhanced both intraradical and extraradical biomass of symbiotic AMF, possibly linked to the suppression of AM-activated defense responses in host roots. Unexpectedly, exogenous myristate disrupted the C-P exchange in AM symbiosis, resulting in reduced mycorrhizal P benefits for plants and decreased allocation of symbiotic C to AMF, indicating that the application of exogenous myristate may not be a promising strategy to enhance AM symbiosis. These findings provide new insights into the nutritional interactions in AM symbiosis, which are crucial for understanding and managing this ancient relationship in a changing world.

**Fig. 8.**
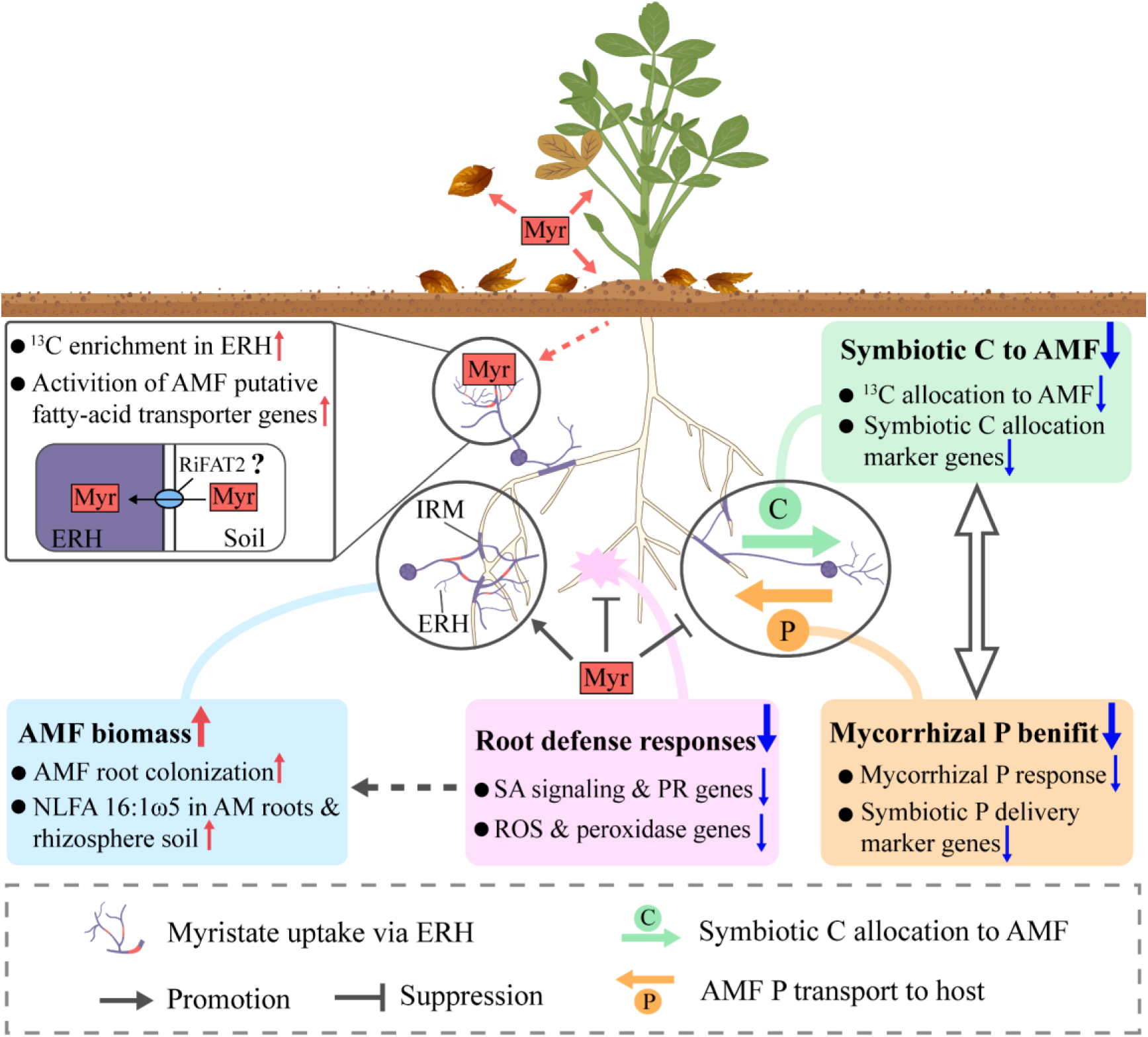
A proposed model for direct uptake of exogenous myristate by symbiotic arbuscular mycorrhizal fungi (AMF) and its regulatory effects on the arbuscular mycorrhizal symbiosis. Symbiotic AMF can uptake exogenous myristate, which is frequently found in diverse soil and plant environments, possibly through the fungal fatty acid transporters RiFAT2 in the extraradical hyphae (ERH), as indicated by ^13^C enrichment and transcriptional activation of fatty acid transport and metabolism genes in AMF ERH. Exogenous myristate increased both intraradical and extraradical fungal biomass, possibly linked to the suppression of AM-activated defense responses in host roots. Unexpectedly, as a non-symbiotic carbon source, exogenous myristate disrupted the carbon (C)-phosphorus (P) exchange in AM symbiosis, resulting in reduced mycorrhizal P benefits for plants and decreased allocation of symbiotic C to AMF. Question mark or dashed arrow indicates putative or unclear mechanisms. IRM, intraradical mycelium; SA, salicylic acid; ROS, reactive oxygen species; PR, pathogenesis-related; NLFA 16:1ω5, neutral lipid fatty acid 16:1ω5 (AMF marker).

## MATERIALS AND METHODS

### Biological materials and experimental design

*In vitro* cultures of AMF species *Rhizophagus irregularis* (strain: MUCL 41833), *R. intraradices* (strain: MUCL 49410), and *R. diaphanus* (strain: MUCL 49416) associated with carrot (*Daucus carota*) hairy roots were generously supplied by the Glomeromycota *In vitro* Collection (GINCO).

Our study consisted of three interconnected and complementary experiments. In brief, we investigated whether exogenous myristate could be directly absorbed by *R. irregularis*, *R. intraradices*, and *R. diaphanus* (Exp. 1), and examined its effect on the growth and development of *R. irregularis* (Exp. 2), using Ri T-DNA*-*transformed carrot roots as the AMF host. In Exp. 3, a field survey was conducted to evaluate the environmental distribution of myristate in soil and plants, along with a pot experiment evaluating the effect of exogenous myristate on C-P nutritional exchanges in AM symbiosis between *R. irregularis* and two host species.

### Exp. 1: ^13^C_1_-labeled myristate (^13^C_1_-Myr) tracing in two-compartment AMF-hairy root co-culture systems

In Exp. 1, ^13^C_1_-Myr tracing experiments were performed to determine whether exogenous myristate can be directly absorbed by the extraradical hyphae of *Rhizophagus irregularis* (two independent trials), *R. intraradices*, and *R. diaphanus* associated with carrot hairy root cultivated in solid modified Strullu–Romand (MSR) medium, utilizing a two-compartment Petri dish system (90 × 15 mm, see Fig. 1a) as described by St-Arnaud et al. (1996), with minor modifications (see Supporting Information Methods S1).

When dense AMF extraradical hyphae were observed in the hyphae compartment (HC), 12–14 weeks later, 200 μL of 0.1 mM myristate (ZZBIO, China, https://www.zzsrm.cn/product/289111^)^ with 98% ^13^C_1_ labeling (^13^C_1_-Myr) or unlabeled (^12^C-Myr, control) myristate (Sigma-Aldrich, China, https://www.sigmaaldrich.cn/CN/zh/product/sigma/m3128) was added weekly to the HC until harvest, two or three weeks later. At harvest, AMF extraradical hyphae from the HC and root compartment (RC) were collected separately by dissolving the medium in 10 mM sodium citrate, filtering through a 37 µm nylon mesh, and thoroughly rinsing five times with sodium citrate followed by ultrapure water. To check for possible contamination of ^13^C-myristate from the HC to the RC, a small portion of the MSR medium from the RC was also collected randomly from three ^13^C_1_-Myr and three ^12^C-Myr (control) treatments, with roots and AMF hyphae removed using tweezers under a stereomicroscope (M250FA, Leica, Germany), and then used for determination of ^13^C enrichment. During the experiment, the MSR medium in the RC was not contaminated by ^13^C-myristate from the HC, as evidenced by the identical ^13^C/^12^C ratios of the MSR medium collected in the RC in ^13^C_1_-Myr (1.0966 ± 0.0012) and ^12^C-Myr (1.0967 ± 0.0032) treatments.

The determination of ^13^C enrichment in AMF extraradical hyphae and solid MSR medium was performed as previously described (Bao et al., 2019), with the details provided in the Supporting Information Methods S2.

### Quantitative real-time PCR (qRT-PCR) targeting fungal marker genes and quantification of AMF colonization intensity (Exp. 1)

In Exp. 1, the expression of fungal marker genes related to fatty acid uptake (*RiFAT1* and *RiFAT2*, Brands and Dörmann, 2022) and metabolism—β-oxidation (*fatty acid degradation 1* (*RiFAD1*)), palmitvaccenic acid synthesis (*oleate desaturase-like enzyme 1* (*RiOLE1*), Brands et al., 2020), and the TCA cycle (*citrate synthase 1* (*RiCIT1*))—in extraradical hyphae of *R. irregularis* from the HC, with or without myristate addition, was assessed, using the elongation factor-1 beta gene (*RiEF-1β*) as the reference gene (Das et al., 2022). Relative gene expression levels were calculated using the comparative 2^−ΔΔCt^ method (Livak and Schmittgen, 2001), with four independent biological replicates and three technical replicates per biological replicate. The collection of extraradical hyphae from the HC and qRT-PCR procedures is detailed in the Supporting Information (Method S3, Table S1).

Carrot hairy roots in the RC from the two-compartment AMF-hairy root co-culture system (Exp. 1) were also harvested using sterile tweezers and used to assess AMF colonization intensity (see Supporting Information Methods S4).

### Exp. 2: Effects of exogenous myristate on AMF development in single-compartment AMF-hairy root co-culture systems

In Exp. 2, we assessed the effects of exogenous myristate at varying concentrations on spore germination and the development of *R. irregularis* intraradical and extraradical structures associated with carrot hairy roots in a single-compartment co-culture system.

Briefly, three 4-cm-long sterile carrot hairy root segments were cultivated in MSR medium with or without *R. irregularis* inoculation (200 sterilized spores per Petri dish). Potassium myristate solution (0, 0.1, 0.5, 1, 5, or 10 mM, 200 μL) was applied to the plates weekly until harvest, with five biological replicates (plates) per treatment. The germination rate of AMF spores, as well as extraradical hyphal branching and length, were monitored under a stereomicroscope (M250FA, Leica, Germany) and calculated using ImageJ (version 2.0.0) at 4 days post-inoculation (Schneider et al., 2012). Spore density was monitored every 2 weeks starting 8 weeks after AMF inoculation.

Carrot hairy roots and MSR medium were harvested at 14 weeks post-AMF inoculation. The roots and solid culture medium were carefully separated using sterile tweezers. The medium was weighed and dissolved in 10 mM sodium citrate, and the AMF collection was preliminarily separated into hyphae and spores using a blender, followed by filtration through a 250 µm and 37 µm nylon meshes, respectively, and manual selection with tweezers under a stereomicroscope (M250FA, Leica, Germany). Biomasses of the extraradical hyphae and spores were oven-dried at 80°C for 48 hours and quantified separately. To evaluate the development of AMF intraradical structures in the carrot hairy roots, hyphal, vesicle, and arbuscular colonization intensities were assessed (Supporting Information Methods S4).

### Exp. 3: Environmental distribution of myristate and the effect of exogenous myristate on AM symbiosis

In Exp. 3, a field survey and a pot experiment were conducted. The field survey aimed to evaluate the environmental distribution of myristate in soil and plants. We analyzed myristate concentrations in bulk soil (0-10 cm) and in plant leaves, roots, leaf litter, and rhizosphere soil from the dominant species *O. sativa*, *Axonopus compressus*, and *Morus alba* in a paddy field, grassland, and woodland, respectively. The procedures regarding sample collection, pretreatment, and analysis of myristate were detailed in Supporting Information Methods S5.

The pot experiment in Exp. 3 was conducted to assess the effect of exogenous myristate on C-P nutritional exchanges in AM symbiosis between *R. irregularis* and two phylogenetically distant plant species—the AM-responsive plant *Medicago sativa* L. cv. Sanditi (alfalfa, dicot) and the AM-nonresponsive plant *Oryza sativa* L. cv.

Zhonghua 11 (rice, monocot) (Hata et al., 2010; Li et al., 2024). This was achieved using ^13^C-neutral lipid fatty acid (NLFA)-isotope ratio mass spectrometry technology, along with physiological and transcriptome analyses. The pot experiment employed a full factorial design with four treatment factors: myristate addition (with or without 0.1 mM myristate), AMF inoculation (with or without *R. irregularis*), host species (alfalfa or rice), and soil P level (relatively low or high). Ten biological replicates were included per treatment, totaling 160 pots. The experimental procedures for plant cultivation, P or myristate treatment, plant harvest, and measurement of plant P concentration are detailed in Supporting Information Methods S6.

### RNA sequencing (RNA-seq) on alfalfa and rice roots and bioinformatics (Exp. 3)

In Exp. 3, alfalfa and rice roots inoculated with AMF, with or without myristate application at two P levels, were subjected to RNA sequencing (RNA-seq). Three biological replicates (roots from a single pot per replicate) were randomly selected from ten replicates per treatment, resulting in a total of 24 samples. The procedures for RNA-seq and bioinformatics followed our previous study (Wang et al., 2023), with details provided in Supporting Information Methods S7.

### Quantitative real-time PCR (qRT-PCR, Exp. 3)

In Exp. 3, RNA extracted from alfalfa or rice roots was reverse-transcribed into cDNA using the QuantiTect Reverse Transcription Kit (Qiagen, Valencia, CA). qRT-PCR was performed, with three independent biological replicates (randomly selected from ten replicates) for each treatment and three technical replicates per biological replicate. The relative expression levels of plant and fungal genes involved in C delivery from plants to AMF (*MsSTR2*, *OsSTR2*, *RiFAT1*, *RiFAT2*, and *RiMST2*) (Brands and Dörmann, 2022; Das et al., 2022), the mycorrhizal P pathway (*MsPT8*, *OsPT11*, and *RiPT*) (Yang et al., 2012; Fiorilli et al., 2013), and AM-activated plant defense—specifically, reactive oxygen species balance (*MsAOX3*), stress responses (*MsUGT85A*, *MsPRXP7*, and *OsPRX39*), and typical defense marker (*OsCHT1* and *OsTGA2*) (Rehman et al., 2018; Suleman et al., 2021; Zhang et al., 2021)—were analyzed to characterize the impact of exogenous myristate on C-P nutritional exchange in AM symbiosis and the underlying mechanisms at the transcriptional level. The alfalfa *actin 2* (*MsACT2*), rice *cyclin 2* (*OsCyc2*), or *R. irregularis elongation factor-1 beta* (*RiEF-1β*) genes were used as reference genes (Yang et al., 2012; Das et al., 2022), with the procedures and primers provided in the Supporting Information (Method S3, Table S1).

### ^13^C labeling of plants, lipid extraction, measurement of AMF signature fatty acids, and quantification of 13C enrichment and 13C flow (Exp. 3)

In Exp. 3, ^13^C labeling of plants, lipid extraction, measurement of AMF signature fatty acids (NLFA 16:1ω5, Olsson et al., 2005), and quantification of ^13^C enrichment and ^13^C flow to NLFA 16:1ω5 were performed as described in previous studies (Olsson et al., 2005; Bao et al., 2019), with details provided in Supporting Information Methods S8.

## Statistical analysis

Mycorrhizal response was calculated based on plant biomass (mycorrhizal growth response) and plant P concentration (mycorrhizal P response), respectively, following Corrêa et al. (2014): Mycorrhizal response (%) = (AM − NM) / NM × 100%, where AM denotes the observed value for the target parameter (biomass or P concentration) in mycorrhizal plants, and NM represents the mean value of nonmycorrhizal plants. Statistical analyses were performed using one-way ANOVA to compare ^13^C enrichment levels in ERH among different AMF species and to evaluate the effect of myristate on AMF growth. Two-way ANOVA was used to evaluate the effects of myristate, phosphorus level, and their interaction on AMF colonization intensity, the concentration of AMF signature fatty acids, the expression levels of *RiEF-1β*, and metrics of plant carbon-phosphorus exchange. A three-way ANOVA was applied to evaluate the effects of AMF inoculation, myristate, phosphorus level, and their interactions on plant phosphorus concentration. Post-hoc comparisons were conducted using Tukey’s test (equal variances) or Games-Howell’s test (unequal variances). Significant differences between AM and NM groups or between myristate-treated and untreated groups were determined using Student’s t test (for normally distributed data) or Mann-Whitney U-test (for non-normally distributed data).

## Author Contributions

**Hanwen Chen:** Investigation, Formal analysis, Methodology, Visualization. **Tian Xiong:** Investigation, Formal analysis, Data curation, Methodology. **Baoxing Guan:** Writing – original draft, Investigation, Formal analysis, Data curation, Visualization. **Jiaqi Huang:** Investigation, Methodology, Resources. **Danrui Zhao:** Investigation, Resources. **Yao Chen:** Investigation, Resources. **Haoran Liang:** Investigation, Visualization. **Yingwei Li:** Investigation, Resources. **Jingwen Wu:** Investigation, Resources. **Shaoping Ye:** Resources. **Ting Li:** Resources. **Wensheng Shu:** Supervision. **Jin-tian Li:** Supervision. **Yutao Wang:** Conceptualization, Data curation, Writing – original draft, Writing – review & editing, Funding acquisition, Supervision, Project administration.

## Supporting information

Source data for figures

## Acknowledgments

We thank the financial support from the National Key R&D Program of China (Key special Project for Marine Environmental Security and Sustainable Development of Coral Reefs 2022-702), the National Natural Science Foundation of China (nos. 32272203 and 31772397), the Fujian Province Science and Technology Plan Project (no. 2024I1011), the Shenzhen Science and Technology Program (KQTD20190929172839621), the Agricultural and Social Development Project of Guangzhou Municipal Science and Technology Bureau (no. 202206010058), and the Guangdong Provincial Key Laboratory of Plant Resources (no. 2023PlantKFP03).

## Data Availability

The RNA-seq data generated in this study have been deposited in the Sequence Read Archive at the National Center for Biotechnology Information under BioProject ID PRJNA1083641. The raw data are provided as a Source Data File.

