## Supporting Information for "Exogenous myristate fuels the growth of symbiotic arbuscular mycorrhizal fungi but disrupts their carbon-phosphorus exchange with host plants"

**The following Supporting Information is available for this article:**

**Fig. S1** Effects of exogenous myristate on the abundance of hyphae and vesicles of arbuscular mycorrhizal fungi (AMF, *Rhizophagus irregularis*) in the roots of *Medicago sativa* and *Oryza sativa*.

**Fig. S2** Effects of exogenous myristate on plant biomass and mycorrhizal growth responses (MGRs) in *Medicago sativa* and *Oryza sativa*.

**Fig. S3** Overall transcriptomic responses of arbuscular mycorrhizal fungi (AMF)-colonized roots of *Medicago sativa* and *Oryza sativa* to exogenous myristate under low (LP) and high phosphorus (HP) conditions.

**Fig. S4** Effects of exogenous myristate on the expression of genes associated with stress or defense gene ontology (GO) terms in response to arbuscular mycorrhizal fungi colonization in **(a)** *Medicago sativa* and **(b)** *Oryza sativa* roots under low (LP) and high phosphorus (HP) conditions.

**Table S1** Primers used for quantitative real-time PCR targeting genes involved in carbon-phosphorus exchange in arbuscular mycorrhiza or fungal fatty acid metabolism.

**Methods S1** Construction and maintenance of the two-compartment Petri dish system used in Exp. 1

**Methods S2** Determination of $^{13}C$ enrichment in arbuscular mycorrhizal fungi (AMF) extraradical hyphae and solid medium in Exp. 1

**Method S3** Quantitative real-time PCR (qRT-PCR) targeting fungal marker genes in Exp. 1

**Methods S4** Assessment of arbuscular mycorrhizal fungi (AMF) colonization intensity in roots (Exps. 1, 2, and 3)

**Method S5** Evaluation of the environmental distribution of myristate in soil and plant environments in the field survey in Exp. 3

**Methods S6** Plant cultivation, P or myristate treatment, plant harvest and measurement of plant P concentration in Exp. 3

**Methods S7** RNA extraction, Illumina sequencing, and bioinformatics for alfalfa and rice roots in Exp. 3

**Methods S8** $^{13}C$ labeling of plants, lipid extraction, measurement of arbuscular mycorrhizal fungi (AMF) signature fatty acids, and quantification of $^{13}C$ enrichment and $^{13}C$ flow (Exp. 3)

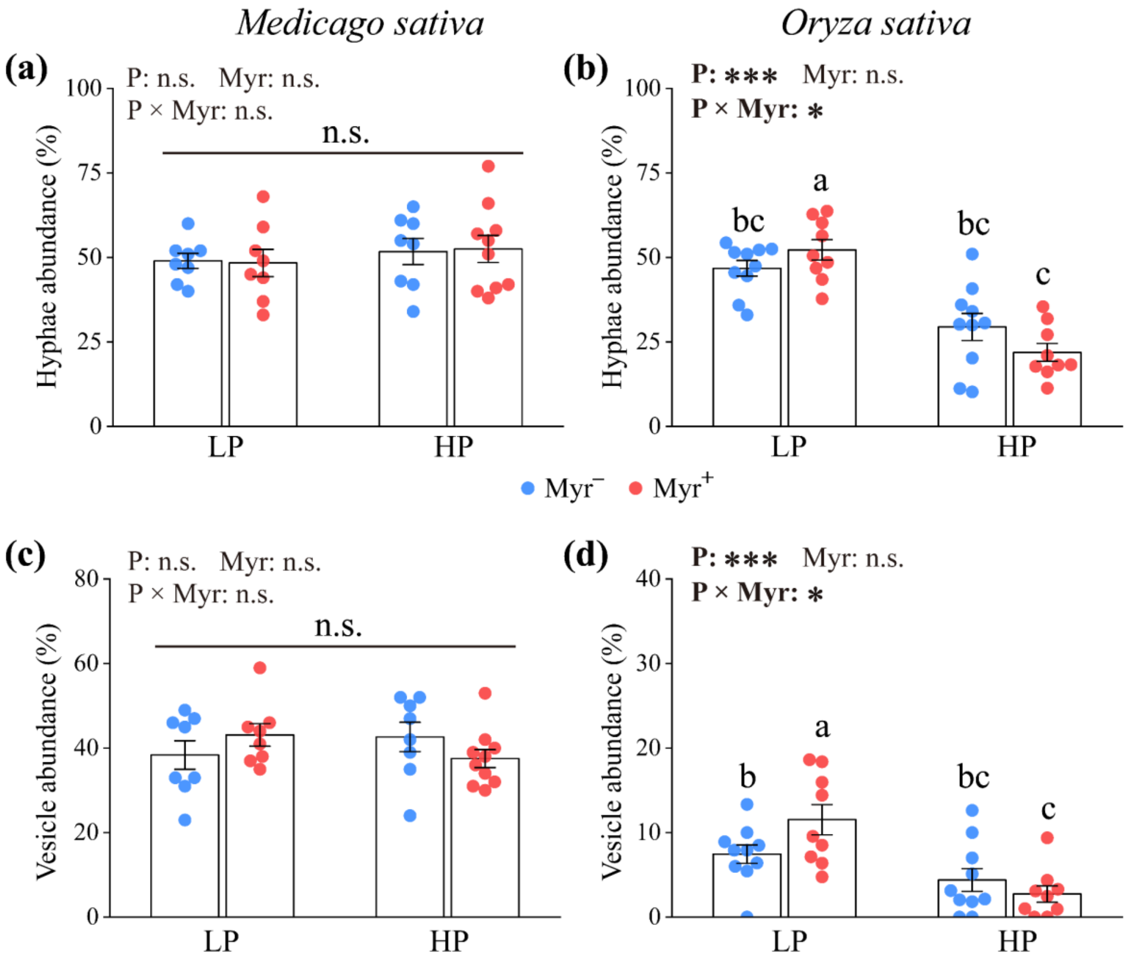


**Fig. S1** Effects of exogenous myristate on the abundance of hyphae and vesicles of arbuscular mycorrhizal fungi (AMF, *Rhizophagus irregularis*) in the roots of *Medicago sativa* and *Oryza sativa*. **(a, b)** AMF hyphae and **(c, d)** vesicle abundance in the roots of *M. sativa* and *O. sativa* treated with 0.1 mM myristate ($Myr^+$) or $ddH_2O$ ($Myr^-$) under low (LP) or high (HP) phosphorus conditions. Means ± SE (n = 8−10); different letters indicate significant differences at $P < 0.05$ (one-way ANOVA followed by Tukey's post hoc test).

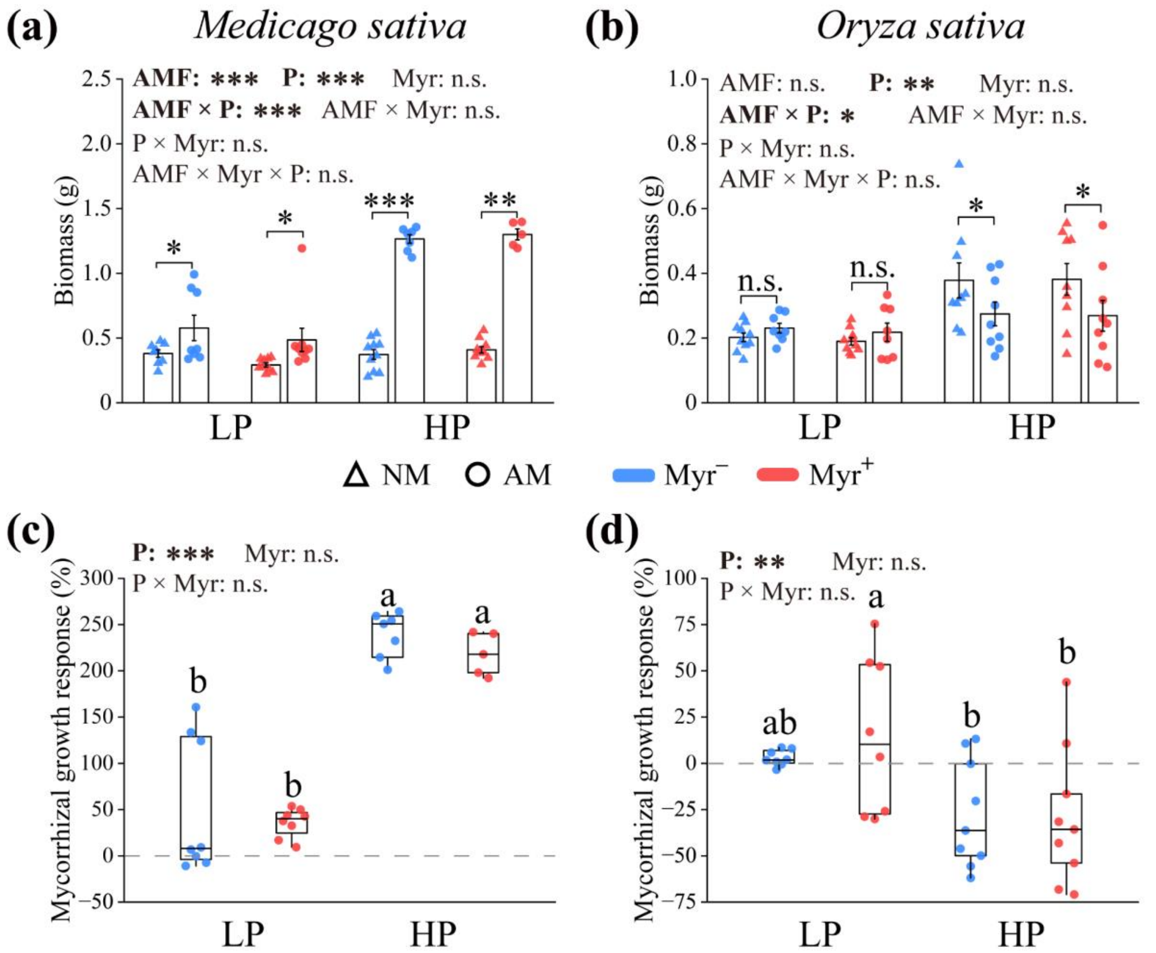


**Fig. S2** Effects of exogenous myristate on plant biomass and mycorrhizal growth responses (MGR) in *Medicago sativa* and *Oryza sativa*. **(a, b)** Biomass and **(c, d)** MGR of *M. sativa* and *O. sativa* were measured following treatment with 0.1 mM myristate (Myr$^{+}$) or dd$H_2O$ (Myr$^{-}$) under low (LP) and high (HP) phosphorus conditions. Bar plots show means ± SE; Box plots display the median (center line), interquartile range (box bounds), and the largest (top whisker) and smallest (bottom whisker) values within 1.5 × interquartile range (n = 5−10). Different letters indicate significant differences at $P < 0.05$ (one-way ANOVA followed by Tukey's post hoc test). *, **, and *** represent statistical significance at the 0.05, 0.01, and 0.001 probability levels (Student's *t*-test), respectively.

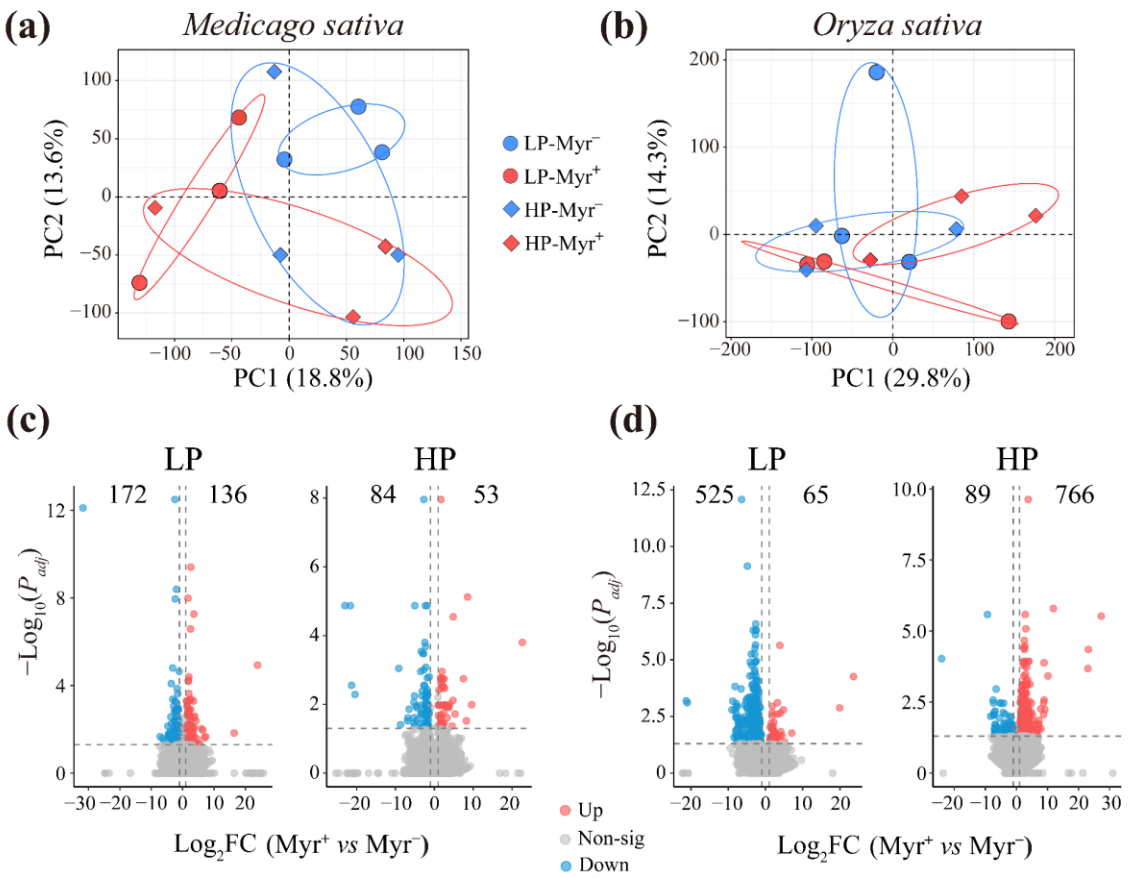


**Fig. S3** Overall transcriptomic responses of arbuscular mycorrhizal fungi (AMF)-colonized roots of *Medicago sativa* and *Oryza sativa* to exogenous myristate under low (LP) and high phosphorus (HP) conditions. **(a, b)** Principal component (PC) analysis of gene expression transcripts per million (TPM) values; **(c, d)** Volcano plot depicting the numbers of up- and downregulated differentially expressed genes (DEGs) between AMF-colonized roots treated with 0.1 mM myristate ($Myr^+$) or dd$H_2O$ ($Myr^-$) under low (LP) and high (HP) phosphorus conditions. Genes (transcripts) with an adjusted *P*-value < 0.05 and $|\log_2(\text{fold change})| > 1$ identified by DESeq2 were considered as differentially expressed.

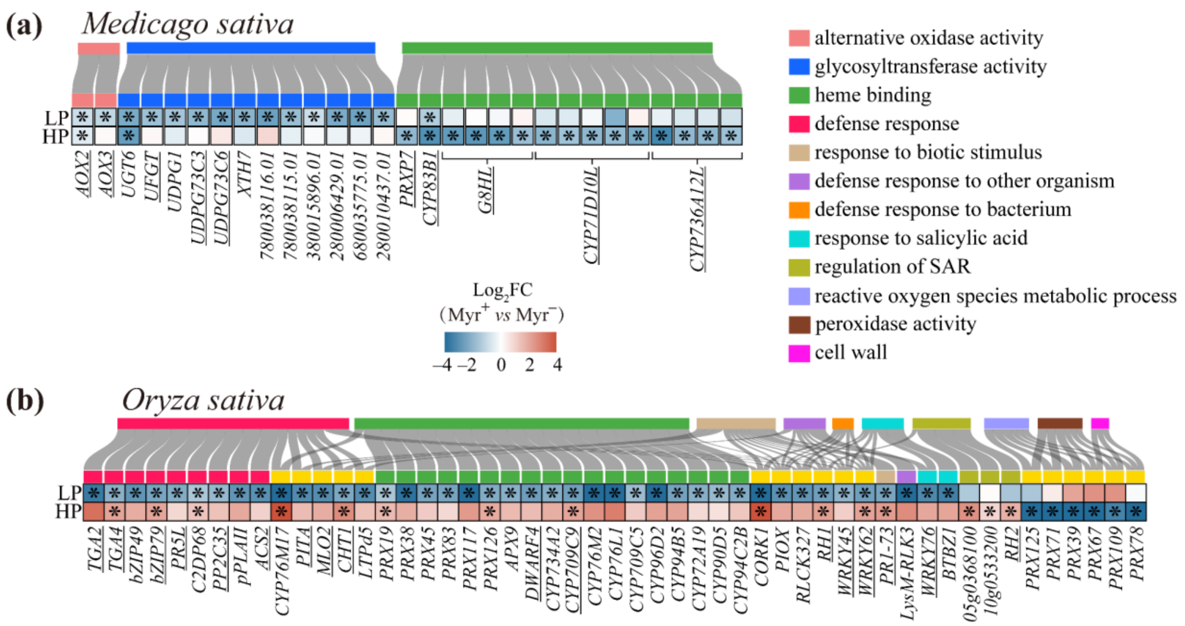


**Fig. S4** Effects of exogenous myristate on the expression of genes associated with stress or defense gene ontology (GO) terms in response to arbuscular mycorrhizal fungi colonization in **(a)** *Medicago sativa* and **(b)** *Oryza sativa* roots under low (LP) and high phosphorus (HP) conditions. Underlined are established stress- or defense-related transcription factors or marker genes. Genes associated with multiple GO terms are highlighted in yellow. * indicates statistical significance between myristate ($Myr^+$) and dd-$H_2O$ ($Myr^-$) treatments at the 0.05 probability level (Benjamini-Hochberg method). Source data are provided as a Source Data file.

**Table S1** Primers used for quantitative real-time PCR targeting genes involved in carbon-phosphorus exchange in arbuscular mycorrhiza or fungal fatty acid metabolism

| Primer | Forward sequence | Reverse sequence |
|---|---|---|
| ***Medicago sativa*** | | |
| MsACT2 | CACGAGACCACCTACAACTCTATC | CATACGGTCAGCAATACCTGG |
| MsPT8 | CTTCACTGATGCCTATGATCTGTTCT | TCCGAGCCAGCCGAATACTAAT |
| MsSTR2 | GCCTAATCACTCACATGCCGTT | CGCTGCTCAAGAAGTATCCACAA |
| MsAOX3 | AGGATGATGTCTTCGCAAGC | TTCTTCTCCGTCGATTTCGC |
| MsUGT85A | GAGCTAGAGTGTTTGGAATGGC | TAGCAAGTCCCCAAGCAAAC |
| MsPRXP7 | ATTCCTGGACCATCTTCTGACC | TGACACTCAGTTTGCCCTATGG |
| ***Oryza sativa*** | | |
| OsCyc2 | GACGAGGTGTTCAAGTTCAAGC | ATGAAGAACTGGGACCCGTTAG |
| OsPT11 | TTCGGCATTCAGAAGCTCAC | AGAAGAAGCCGAGCATGTTG |
| OsSTR2 | ATTTCCACGATGAAGTACCCTTAC | GGTAGAAGAGGACACGGTAGAAGA |
| OsCHT1 | TTCGACCAGATGCTTCTCCAC | AAGGCGTCGTAGGTGTAGAAG |
| OsTGA2 | GCTCTCTTCCAACACCGAAG | GAGTGGAACTTGCTGCCATC |
| OsPRX39 | TCCACTTCCACGACTGCTTC | AACTTCTCCGACACGATGCTC |
| ***Rhizophagus irregularis*** | | |
| RiEF-1β | CCCATGCAGCTCGATGGTA | TGCCAGGAAGTGAAGAAAATGA |
| RiPT | GGTGTCGGTATTGGAGGAGA | GAGGACAGCGAAACCCATTA |
| RiFAT1 | TTATGCGGCTTTACTCCGT | ATGATTTACCACCAGGATGTTC |
| RiFAT2 | TATACGCACGATTGGGATCC | TGAGGGTGCGTATATGGATC |
| RiMST2 | GTTAATGGTCTTGTCAATATGTTAG | AAATGTTTTCCCAACGATTCATCA |
| RiFAD1 | TGGTATTCATTCGTGCCATGA | GCAGCGCGTGCAAGTG |
| RiOLE1 | CAACGGGAAACGACCCAAAT | GTCTTGGTGACATACGGAATG |
| RiCIT1 | CTACATGGCCTTGCTAATCAAGAA | CACCAATCGCGTCTCTCATCT |

**Methods S1 Construction and maintenance of the two-compartment Petri dish system used in Exp. 1**

The Petri dish was divided into two compartments by a central plastic barrier. One compartment (root compartment, RC) contained 25 mL of modified Strullu–Romand (MSR) solid medium for the growth of carrot hairy roots and AMF hyphae. The other compartment (hyphal compartment, HC), into which only AMF hyphae, and not roots, were allowed to grow, contained 15 mL of solid MSR medium separated from the solid medium in the RC. Five 4-cm-long sterile carrot hairy root segments were initially co-cultivated with 200 AMF spores in the RC in an incubator at 27°C in the dark. The Petri dishes were positioned at an angle of approximately 45 degrees (with the HC underneath) in the incubator to facilitate hyphal growth from the RC into the HC and to prevent contamination of exogenous $^{13}$C-myristate added in the HC from reaching the RC. The roots were routinely trimmed as needed to prevent them from growing into the HC.

**Methods S2 Determination of $^{13}C$ enrichment in arbuscular mycorrhizal fungi (AMF) extraradical hyphae and solid medium in Exp. 1**

AMF hyphae collected from the hyphal compartment (HC) or root compartment (RC), as well as modified Strullu–Romand (MSR) solid medium from the RC, were sealed in tin capsules and analyzed using an elemental analyzer coupled to an IRMS (EA-IRMS, Flash EA 2000 and Delta V Plus; Thermo Fisher Scientific, Bremen, Germany). The $^{13}C$:$^{12}C$ ratio of $CO_2$ from combusted samples was measured with a precision of ±0.2‰ and expressed as atom ‰ $^{13}C$, using a sucrose standard calibrated against the Vienna Pee Dee Belemnite standard. The $^{13}C$ enrichment in AMF extraradical hyphae from the HC and RC under $^{13}C$-Myr treatment was calculated by subtracting the background $^{13}C$ levels determined from non-labeled ($^{12}C$-Myr) extraradical hyphae in the corresponding HC and RC, respectively.

**Method S3 Quantitative real-time PCR (qRT-PCR) targeting fungal marker genes in Exp. 1**

In the two-compartment AMF-hairy root co-culture system (Exp. 1), extraradical hyphae of *R. irregularis* collected from the HC were thoroughly washed, flash-frozen in liquid nitrogen, and stored at −80°C prior to RNA extraction. Total RNA was extracted from 30–50 mg of extraradical hyphae using the Plant RNeasy® Kit (Qiagen, Hilden, Germany) according to the manufacturer's instructions. Extracted RNA was reverse-transcribed into cDNA using the QuantiTect Reverse Transcription Kit (Qiagen, Valencia, CA). qRT-PCR was performed on a QuantStudio™ 5 Real-Time PCR System (Applied Biosystems, CA, USA) with the SYBR ROX Mastermix Kit (Qiagen, Valencia, CA), following the manufacturer's protocol. Relative gene expression levels were calculated using the comparative $2^{-\Delta\Delta Ct}$ method (Livak and Schmittgen, 2001), with four independent biological replicates and three technical replicates per biological replicate. qRT-PCR primers used are provided in the Supporting Information (Table S1).

**Methods S4 Evaluation of arbuscular mycorrhizal fungi (AMF) colonization intensity in roots (Exps. 1, 2, and 3)**

Briefly, 100 root segments (approximately 1 cm in length) were treated with a 10% KOH solution at 95°C for 10 minutes, decolorized in a 30% $H_2O_2$ solution at 25°C for 3 min, softened in a 1% HCl at 25°C for 3 minutes, and then stained with 0.05% Trypan Blue at 95°C for 15 minutes. The root segments were examined under 400× magnification using a light microscope (Nikon Eclipse 80i, Japan). Colonization rates for hyphae, vesicles, and arbuscules were calculated using the gridline intersect method (Giovannetti and Mosse, 1980).

**Method S5 Assessment of the environmental distribution of myristate in soil and plant environments in the field survey in Exp. 3**

A field survey was conducted to evaluate the environmental distribution of myristate in soil and plants from a paddy field, a grassland, and a woodland in Southern China. We analyzed myristate concentrations in bulk soil (0-10 cm), as well as in plant leaves, roots, leaf litter, and rhizosphere soil from the dominant species *Oryza sativa*, *Axonopus compressus*, and *Morus alba* in the paddy field, grassland, and woodland, respectively.

The paddy field, grassland, and woodland habitats are all located at the Biology Garden of the South China Normal University (23°14′N, 113°35′E). The paddy field is cultivated with local rice varieties and features clay soil with a pH of 5.45. The grassland is a manually landscaped lawn dominated by *A. compressus*, with black sandy loam soil having a pH of 7.40. The woodland consists predominantly of *M. alba* and *Mangifera indica*, with gravelly loam soil that has a pH of 8.29. For sample collection, four sampling sites (replicates) were established for each habitat, with a minimum distance of 5 m between them. In each sampling site, fresh leaves and lateral roots from healthy plants were collected using sterile scissors and shovels. Rhizosphere soil was obtained by brushing the tightly adhered soil from root samples with a sterile brush, after removing loosely attached soil by vigorous shaking. Bulk soil, far from plant roots, was also sampled at a depth of 0-10 cm. Senesced leaf litter from the soil surrounding the sampled plants was collected using forceps and cleaned with a soft brush to remove debris. Before analyzing myristate content, rhizosphere and bulk soil samples were air-dried at room temperature; leaf, root, and leaf litter samples were rinsed with ultrapure water, and surface moisture was carefully absorbed using absorbent paper.

Fatty acids were extracted from 0.2 g of plant or soil samples using a modified Bligh and Dyer method (Bligh and Dyer, 1959). Extracted fatty acids were saponified twice with 5 mL of saturated KOH–MeOH solution for 10 min at 75°C, and then methylated with 5 mL HCl-MeOH solution (5 M) for 10 min at 75°C. Methyl fatty acids were subsequently extracted twice with hexane, evaporated to dryness under nitrogen, and redissolved in 1.5 mL of hexane.

Methyl fatty acids were analyzed using an Agilent 7890B gas chromatograph coupled with a 5975C time-of-flight mass spectrometer (GC-MS, Santa Clara, CA, USA). Separation was performed on a DB-5MS capillary column (5% phenylpolysilphenylene-siloxane; 30 m × 0.25 mm i.d. × 0.25 μm film thickness; Agilent) with high-purity helium (99.999%) as the carrier gas at a constant flow rate of 1.0 mL/min. The oven temperature program was as follows: the initial temperature held at room temperature, increased to 180 °C at 4 °C/min (held for 3 min), then to 280 °C at 2 °C/min (held for 20 min), and finally to 300 °C at 15 °C/min (held for 2 min). The solvent delay was set to 8 min. The mass spectrometer was operated in electron impact mode at 70 eV, scanning from m/z 40 to 700 in full-scan mode. Methyl myristate was identified using MSD ChemStation (version E.02.02.1431, Agilent) by matching its mass spectrum against the NIST14 library. Quantification was based on a calibration curve established with a methyl myristate standard (Macklin, Shanghai, China). The accuracy of the quantification method was confirmed by a spike-recovery test for the methyl myristate standard, with a mean recovery rate of 90.06% $\pm$ 7.83% (n = 3).

**Methods S6 Plant cultivation, P or myristate treatment, plant harvest, and measurement of plant P concentration in Exp. 3**

Alfalfa and rice seeds were germinated for 14 days in sterilized sand before being transplanted into a substrate composed of vermiculite, perlite, and sand (3:1:1, v/v/v). To account for potential biomass differences between the two plant species, three alfalfa seedlings or one rice seedling were planted per pot, containing either 300 or 1000 mL of substrate, respectively, with or without *R. irregularis* inoculation (10 spores/mL of substrate). The plants were grown in a greenhouse under a long-day photoperiod (16 h light/8 h dark) at temperatures ranging from 20–30℃. For myristate application, 5 mL of potassium myristate solution (0.1 mM) or dd$H_2O$ was applied near the plant roots twice a week, which did not affect the growth of rice and alfalfa according to our preliminary experiment. The calculated amount of myristate used in the greenhouse experiment (1.83 mg per pot) corresponds to the myristate content present in roughly 500 g of bulk soil in nature, based on our field survey of myristate levels in Exp. 3. For P treatments, considering differences in soil substrate volumes (300 ml per pot for alfalfa and 1000 ml per pot for rice) and P demand among plant species, 10 mL of Hoagland nutrient solution with either relatively low or high P levels was applied twice weekly: for alfalfa, 25 or 100 μM $KH_2PO_4$ (He et al., 2020), and for rice, 100 or 400 μM $KH_2PO_4$ (Shi et al., 2021). The high N:P ratio observed in alfalfa (low-P treatment: 117.3 ± 75.3 in NM, 31.6 ± 13.9 in AM; high-P treatment: 51.3 ± 16.7 in NM, 28.0 ± 6.0 in AM) and rice (low-P treatment: 39.8 ± 9.3 in NM, 31.2 ± 8.9 in AM; high-P treatment: 32.8 ± 5.8 in NM, 24.0 ± 5.3 in AM) plants at harvest indicated that, regardless of low-P or high-P treatments, plants generally grew under P-limited conditions (Güsewell, 2004), particularly the low-P treated alfalfa, which experienced an extremely P-deficient environment.

All plants were harvested eight weeks after inoculation. They were thoroughly washed with tap water and separated into shoot and root portions. A portion of the fresh root samples was flash-frozen in liquid nitrogen and stored at −80°C for molecular analysis. Another portion of the fresh roots was used to assess AMF colonization intensity following the procedures described in Exp. 1. The remaining plant samples were de-enzymed at 105°C for 30 minutes, dried at 60°C for 7 days, and weighed. Plant P concentration was determined using the molybdenum-antimony colorimetric method.

**Methods S7 RNA extraction, Illumina sequencing, and bioinformatics for alfalfa and rice roots in Exp. 3**

Total RNA was extracted from 100 mg of alfalfa or rice root tissue using the TRIzol Plus RNA Purification Kit (Takara, Dalian, China) following the manufacturer's instructions. Library preparation was conducted with the NebNext Ultra RNA Library Prep Kit (New England Biolabs, Ipswich, MA, USA). Qualified libraries were sequenced on the Illumina HiSeq 4000 platform with 150-bp paired-end reads. Fastp was used for basic statistics and quality filtering of raw data.

Hisat2 (version 2.0.5) was utilized to map the sequencing reads to the rice reference genome (IRGSP-1.0) or the alfalfa reference genome (Zhongmu NO.1, Shen et al., 2020). Salmon (version 1.10.2) quant was applied for gene expression quantification. The DESeq2 R package (version 1.20.0) was utilized to conduct differentially expressed gene (DEG) analysis between plants with ($Myr^+$) and without ($Myr^-$) myristate application (Love et al., 2014). The *P* values obtained were adjusted using the Benjamini-Hochberg correction to control false discovery rate. Genes (transcripts) with an adjusted *P*-value < 0.05 and $|\log_2(\text{fold change})| > 1$ identified by DESeq2 were considered as differentially expressed. The clusterProfiler method was applied for Gene Ontology (GO) enrichment analysis of DEGs (Wu et al., 2021), with GO terms showing an adjusted *P*-value < 0.05 considered significantly enriched.

**Methods S8 $^{13}C$ labeling of plants, lipid extraction, measurement of arbuscular mycorrhizal fungi (AMF) signature fatty acids, and quantification of $^{13}C$ enrichment and $^{13}C$ flow (Exp. 3)**

In Exp. 3, the $^{13}CO_2$ labeling procedure was conducted as described in our previous report, with minor modifications (Bao et al., 2019). Five days before harvest, alfalfa and rice plants (five randomly selected replicates) were placed in a sealed acrylic chamber containing $^{13}CO_2$ and incubated for 4 hours each. The extraction and analysis of signature fatty acids were performed following the method described by Olsson et al. (2005), with slight modifications based on Bao et al. (2019). Fatty acid methyl esters were identified and quantified relative to an added internal standard (fatty acid methyl ester 19:0). NLFA 16:1ω5 was used as a marker for AMF in roots and rhizospheric soil (Olsson and Lekberg, 2022).

The $^{13}C$ enrichment in roots and fatty acid methyl esters was analyzed using IRMS coupled to a Trace GC Ultra gas chromatograph via the ConFlow IV interface (Thermo Fisher Scientific, Bremen, Germany), as previously described (Bao et al., 2019). The chromatographic conditions matched those specified for lipid analysis, using helium (He) as the carrier gas. The effluent from the capillary column passed through an Al tube filled with CuO wires at 860°C. The $\delta^{13}C$ values were calibrated against the V-PDB using the $CO_2$ standard (RM 8542 (IAEA-CH-6, sucrose)) and corrected for the additional non-labeled methyl group introduced during methanolysis. The $^{13}C$ enrichment was calculated by subtracting the background $^{13}C$ levels determined from five non-labeled controls, and the total C flow to NLFA 16:1ω5 was calculated according to: NLFA 16:1ω5-C (μg) × $^{13}C$

enrichment of NLFA 16:1ω5 (%/100) = C flow to NLFA 16:1ω5 (Olsson et al., 2005).
